# p38-MAPK mediated rRNA processing and translation regulation enables PrE differentiation during mouse blastocyst maturation

**DOI:** 10.1101/2020.11.30.403931

**Authors:** Pablo Bora, Lenka Gahurova, Tomáš Mašek, Andrea Hauserova, David Potěšil, Denisa Jansova, Andrej Susor, Zbyněk Zdráhal, Anna Ajduk, Martin Pospíšek, Alexander W. Bruce

## Abstract

**Background:** p38-MAPKs are stress-activated kinases necessary for placental development and nutrient and oxygen transfer during murine post-implantation development. In preimplantation development, p38-MAPK activity is required for blastocyst formation. Additionally, we have previously reported its role in regulating specification of inner cell mass (ICM) towards primitive endoderm (PrE), although a comprehensive mechanistic understanding is currently limited. Adopting live embryo imaging, proteomic and transcriptomic approaches, we report experimental data that directly address this deficit.

**Results:** Chemical inhibition of p38-MAPK activity during blastocyst maturation causes impaired blastocyst cavity expansion, most evident between the third and tenth hours post inhibition onset. We identify an overlapping minimal early blastocyst maturation window of p38-MAPKi inhibition (p38-MAPKi) sensitivity, that is sufficient to impair PrE cell fate by the late blastocyst (E4.5) stage. Comparative proteomic analyses reveal substantial downregulation of ribosomal proteins, the mRNA transcripts of which are also significantly upregulated. Ontological analysis of the differentially expressed transcriptome during this developmental period reveals “translation” related gene transcripts as being most significantly, yet transiently, affected by p38-MAPKi. Moreover, combined assays consistently report concomitant reductions in *de novo* translation that are associated with accumulation of unprocessed rRNA precursors. Using a phosphoproteomic approach, ± p38-MAPKi, we identified *Mybpp1a,* an rRNA transcription and processing regulator gene, as a potential p38-MAPK effector. We report that siRNA mediated clonal knockdown of *Mybpp1a* is associated with significantly diminished PrE contribution. Lastly, we show that defective PrE specification caused by p38-MAPKi (but not MEK/ERK signalling inhibition) can be partially rescued by activating the archetypal mTOR mediated translation regulatory pathway.

**Conclusions:** Activated p38-MAPK controls blastocyst maturation in an early and distinctly transient developmental window by regulating gene functionalities related to translation, that creates a permissive environment for appropriate specification of ICM cell fate.

## Introduction

Following fertilisation, the totipotent mouse zygote proceeds through a series of reductive cleavage divisions and, regulated by polarity dependent differential HIPPO pathway activation, differentiate into a extraembryonic outer epithelium called the trophectoderm (TE – the precursor of the embryonic component of the placenta) or contribute to the blastocyst inner cell mass (ICM)^1,2^. Starting from E3.25, cells of the nascent ICM give rise (by E4.5) to both the pluripotent epiblast (EPI – the progenitor pool of cells of the future foetus, ultimately found deep within the ICM) and the differentiating extraembryonic epithelial primitive endoderm (PrE – a monolayer of polarised superficial ICM cells in contact with the fluid filled cavity) lineages^1^. At the E4.0 stage of this maturation process, the blastocyst no longer occupies a volume equivalent to that of the zygote, and begins a process of non-linear expansion. Under *in vitro* culture conditions this is observed as a period of pulsed expansions and contractions, that eventually culminates in the release of the developing blastocyst from the encapsulating *zona pellucida* and permits uterine implantation and subsequent post-implantation development (E4.5 onwards)^3^.

The appropriate maturation of the mouse blastocyst ICM lineages (E3.25-E4.5) is under the regulation of the fibroblast growth factor-extracellular signal-regulated kinase (FGF4-MEK-ERK) signalling axis and occurs from an initially uncommitted ICM population co-expressing both the pluripotency related transcription factor (TF) NANOG and the PrE-related TF GATA6. Accordingly, FGF4 ligands expressed from specifying EPI progenitors act upon precursor PrE cell populations to destabilise NANOG protein expression and thus enable GATA6 (within the context of activated ERK1/2) to initiate expression of *Sox17* and *Gata4* genes (plus other PrE lineage specific markers, reviewed here^1,2^)^4–9^. This molecular specification of PrE is concomitant with downregulation of GATA6 protein expression in EPI progenitors, resulting in a spatially randomised pattern of mutually exclusive NANOG and GATA6 expression within the ICM by the mid-blastocyst stage (E4.0). This so called ‘salt & pepper’ pattern of EPI and PrE progenitors^10^ is then resolved by a combination of active cell movement, primarily mediated by PrE fated cells migrating towards the cavity, and selective apoptosis of aberrantly segregated cells, resulting in the spatially defined EPI and PrE tissue layers of late blastocyst (E4.5) ICM^11,12^. It has been recently reported that such spatial refinement of the specified ICM lineages positively correlates with expansion of the cavity to facilitate the appropriate development of the ICM^12^. This process is enabled by a metabolically activated TE transporting water (via Aquaporins), sodium ions (via Sodium-Potassium (Na^+^/K^+^) ATPases), chloride ions (via apical Cl^-^ channels) and promoting Sodium-Hydrogen ion Exchange^13–17^.

The p38 family of mitogen activated protein kinases (p38-MAPKs), comprising of p38-α (MAPK14), -β (MAPK11), -γ (MAPK12/ERK6), and -δ (MAPK13/SAPK4), are widely studied stress-activated kinases; particularly in inflammatory disease, but also in development^18–20^. *Mapk14* genetic knockout results in post-implantation mouse embryo lethality, via aberrant placental development causing insufficient oxygen and nutrient transfer^21^. During preimplantation embryo *in vitro* culture, pharmacological inhibition of p38-MAPKs (p38-MAPKi) from the 2-cell stage leads to arrested development around the 8- to 16-cell stage^22^ and inhibition post 16-cell stage has been reported to result in failed TE differentiation and function^23,24^. We have previously demonstrated that p38-MAPKi post-E3.5 (so as not to inhibit TE specification) significantly impairs the differentiation of ICM cells towards the PrE lineage (defined by GATA4 expression); resulting in late blastocyst (E4.5) embryos with increased numbers of uncommitted ICM cells (co-expressing NANOG and GATA6) but minimal effects on EPI cell generation (defined by sole expression of NANOG). Moreover, that this role temporally precedes the appearance of the salt & pepper pattern^25^. Additionally, we have also identified a role for p38-MAPK in mitigating amino acid depletion and associated oxidative stress that is detrimental to blastocyst maturation and the emergence of both ICM cell lineages. Collectively, such observations indicate a potential p38-MAPK related function in the maintenance of metabolic conditions conducive to appropriate PrE differentiation and thus appropriate ICM cell fate derivation during mouse blastocyst maturation. Accordingly, using specific pharmacological inhibitor based interventions, (phospho)proteomic and transcriptomic approaches, we have herein uncovered significant p38-MAPK mediated regulation of protein translation and rRNA processing. This regulation is concomitant with the minimal window of functional p38-MAPK activity (prior to the emergence of the salt & pepper pattern) that we argue is necessary to specifically prime PrE progenitors for differentiation during mouse blastocyst maturation.

## Results

### Inhibition of p38-MAPK activity post E3.5 induces a PrE deficit and blastocoel cavity expansion defect in the developing blastocyst

Employing specific pharmacological ‘inhibition and release’-based experimental regimes (using the compound SB220025), we previously identified p38-MAPK function between E3.5 and E3.75 as necessary for normal derivation of GATA4 expressing PrE cells by the peri-implantation blastocyst (E4.5) stage^25^. Here we find further resolution of the inhibition period to just three hours (E3.5 +4 to +7 hour (h)), is sufficient to significantly impair PrE specification (although in a less robust manner – Fig. 1a (i)); identifying a minimal window of p38-MAPK activity required for appropriate ICM cell fate resolution during blastocyst maturation. In our previous studies, we also observed that p38-MAPKi throughout the entire blastocyst maturation period (E3.5 to E4.5) not only impaired PrE differentiation (recapitulated here, Fig. 1a (ii)) but appeared to cause reductions in blastocyst size and cavity volume, particularly in the absence of exogenous amino acid supplementation^26^. Therefore, as blastocyst cavity expansion and volume have recently been linked to EPI and PrE fate acquistion^12^, we decided to directly quantify late blastocyst cavity volumes following p38-MAPKi throughout the 24 hour maturation period and compare them to control treated embryos (Fig. 1b). On average, we observed significantly smaller cavities upon p38-MAPKi (3316μm^2^) when compared to controls (6255μm^2^). Taken together with our previous observations^25,26^ these data indicate late blastocyst deficient PrE cell phenotypes caused by p38-MAPKi are coincident to significantly reduced cavity volumes.

**Figure 1:**
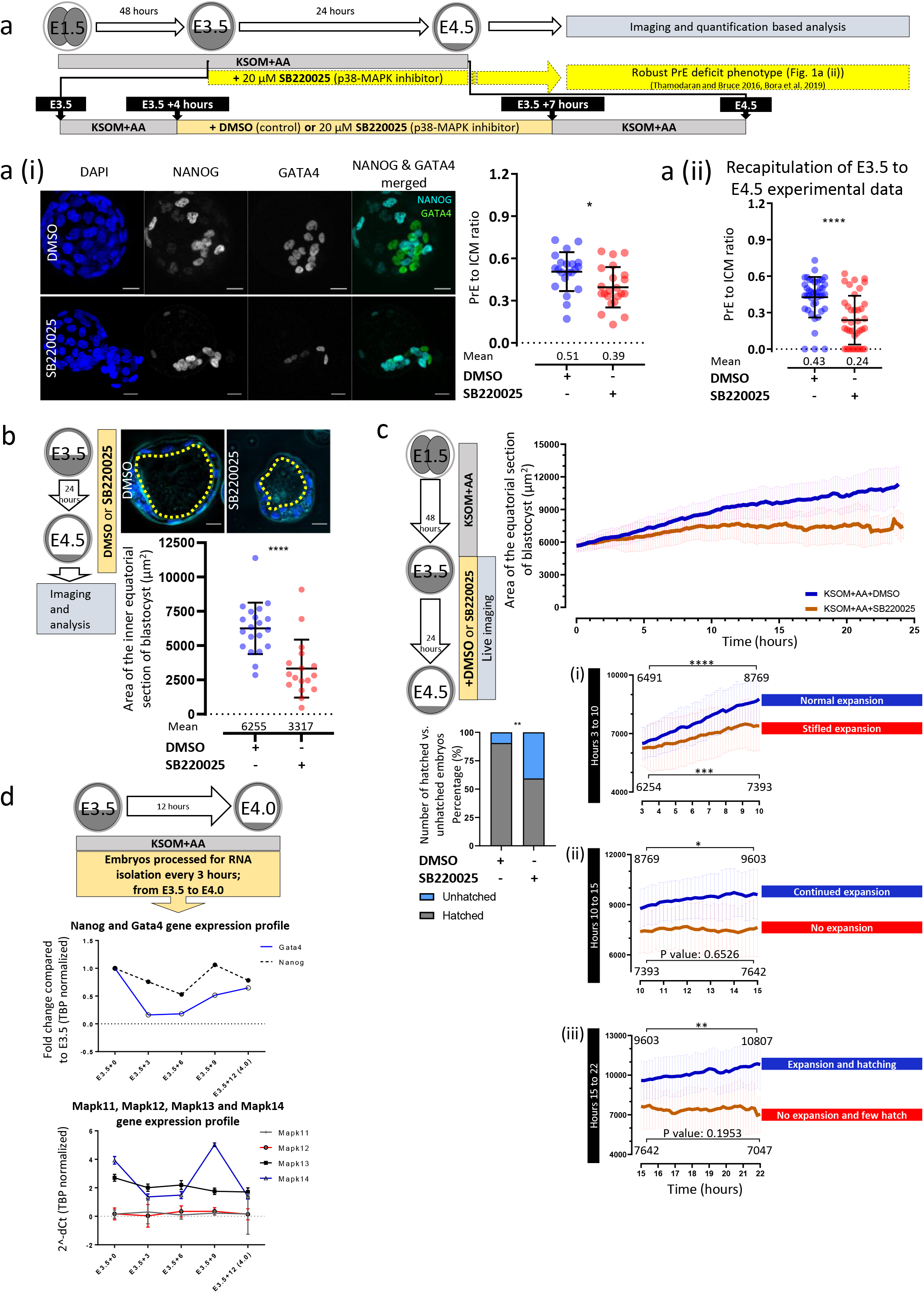
Temporal resolution and morphological effect of inhibiting p38-MAPK signalling in the developing blastocyst. a) Scheme illustrating experimental protocol for resolving the temporal nature of p38-MAPKi in PrE deficit phenotype. Lineage markers NANOG and GATA4 denoting EPI and PrE cells, respectively. a.i Quantification of the ratio of GATA4 positive PrE to entire ICM in control (n=21) and p38-MAPKi (n=22) conditions, in confocal images of E4.5 embryos transiently cultured with DMSO or SB220025 between E3.5 +4 hours (h) to E3.5 +7h (right); means and standard deviation highlighted. Exemplar z-section blastocyst projections shown on left (scale bars = 20μm) a.ii Quantification of the ratio of GATA4 positive PrE to entire ICM in control (DMSO) (n=42) and p38-MAPKi (SB220025) (n=38) conditions on treatment between E3.5 to E4.5 (recapitulation of previously published observations^25,26^); means and standard deviation highlighted. Collated cell count data for each embryo used to calculate the PrE to ICM ratios depicted in a.i and a.ii are provide in supplementary tables S1a and S1b. b) Quantification of cavity equatorial area (μm^2^) in fixed blastocysts cultured (E3.5-E4.5) in control (DMSO; n=20) and p38-MAPKi (SB220025; n=17) conditions. Exemplar confocal equatorial z-sections of each embryo group, with cavity circumference indicated (dashed yellow line) are shown (upper panel – scale bar = 20μm). Collated cavity equatorial area (and volume) quantifications per embryo are provided in supplementary table S1c. c) Equatorial area of control (DMSO; n=32) and p38-MAPKi (SB220025; n=32) blastocysts imaged in live culture from E3.5-E4.5 (supplementary tables S1d). Sub panels segment the recordings between (i) 3h and 10h, (ii) 10h and 15h and (iii) 15h and 22h. Bar graph denotes the percentage of hatching blastocysts observed at the end of the imaging period (E4.5). d) qRT-PCR derived expression levels (normalised to *Tbp)* of *Nanog* and *Gata4* ICM lineage markers (upper) and p38-MAPK paralogous gene mRNAs *(Mapk11/12/13/14* – lower) during early blastocyst maturation (*i.e.* E3.5 +0h, E3.5 +3h, E3.5 +6h, E3.5 +9h and E3.5 +10h).

The *in vitro* imaging of maturing mouse blastocysts (E3.5-E4.5) reveals phases of pulse-like oscillations in blastocyst and consequently blastocoel cavity volume expansion and contraction, that in the clinic are often used as reporters of blastocyst viability in assisted reproductive technologies^27,28^. Having established reduced blastocyst cavity volume as a phenotype of p38-MAPKi in fixed embryos, we asked if the phenotype is marked by a failure to undergo such pulse-like oscillations. Live time-lapse recordings revealed that blastocysts developing under p38-MAPKi actually did expand and pulse akin to control blastocysts during the first three hours of culture (Fig. 1c and supplementary videos). However, in the subsequent seven hours the rate of expansion in the inhibited blastocysts was significantly slower (expanding from an average equatorial area of 6254μm^2^ to 7393μm^2^) than that observed in the control group (increasing from 6491μm^2^ to 8769μm^2^ – Fig. 1c (i)). Between the tenth and fifteenth hours, control embryos continued expanding but at a much reduced rate (achieving an area of 9603μm^2^), whereas the p38-MAPKi blastocysts did not significantly expand (with an average area of 7642μm^2^ – (Fig. 1c (ii)). From the fifteenth hour to the end of the imaging period *(i.e.* after 24 hours and at E4.5), the control embryos continued growing at the slower rate (culminating in an average size of 10807μm^2^) and 90.6% hatched from the *zona pellucida,* whereas the p38-MAPKi embryos remained unexpanded (averaging a size of 7047μm^2^) and in 40.6% of cases failed to hatch, often associated with cavity collapse (Fig 1c & 1c (iii)). In summary, these data reveal p38-MAPKi blastocysts can initially expand their cavities in a manner indistinguishable from controls but after 3 hours this rate of expansion decreases and eventually ceases after 10 hours, resulting in smaller blastocyst less able to hatch.

We next assayed the mRNA gene expression level of each of the four p38-MAPK paralogs, *Mapk11, Mapk12, Mapk13* and *Mapk14* (plus the ICM lineage markers, *Nanog* and *Gata4)* across the identified early blastocyst maturation window most sensitive to p38-MAPKi *(i.e.* from E3.5 and every three hours thereafter until E4.0 – note, our previous study indicates p38-MAPKi after E4.0 does not result in any PrE differentiation phenotypes^25^). This analysis revealed *Mapk13* and *Mapk14* (encoding the p38-δ and -α isoforms, respectively) as the only robustly expressed p38-MAPK gene paralogs at this time. Moreover, we observed increasing *Nanog* and *Gata4* transcript levels between the E3.5 +6h and E3.5 +9h time-points (related to ICM cell fate derivation) that coincided with an increase in *Mapk14* mRNA expression (Fig. 1d). Therefore, given this developmental period partly coincides with the minimal window of p38-MAPKi sensitivity in regard to PrE specification (Fig. 1a) and the transition from impaired to blocked cavity expansion (Fig. 1c), we reason the p38-MAPK isoform expressed from the *Mapk14* gene paralog as the best candidate mediating the phenotypes observed. This is substantiated by the fact the *Mapk14* genetic knockout alone is embryonic lethal^21,29^ (unlike the three other paralogs^30,31^).

Collectively, these introductory data (together with our previous reports^25,26^) confirm p38-MAPK (most probably the p38-α isoform) has a transient role during early blastocyst maturation, that feeds forward to ensure appropriate specification of PrE cells. Specifically, the developmental period of most significance is between third and tenth hours after E3.5 (and can be refined to a minimal 3-hour window within which p38-MAPKi is sufficient to significantly impair PrE differentiation).

### p38-MAPK inhibition in developing blastocysts affects the translation machinery

p38-MAPKs are known to effect many different pathways and physiological states^18,19,32^. However, to our knowledge, there have been no global analyses of the (phospho)proteomic changes brought about by p38-MAPK activity in developing mammalian blastocysts. Therefore, using triplicate samples of 300 mouse blastocysts per condition and a mass-spectrometry read out, we assayed both the differential proteome and phosphoproteome induced by 7 hours of p38-MAPKi during early blastocyst maturation (*i.e.* between E3.5 +2 and +9h – informed from our introductory experiments).

Accordingly, we detected 77 proteins significantly downregulated and 53 upregulated after p38-MAPKi, as compared to the control group (supplementary table S2b and Fig. 2c). A subsequent ontological analysis revealed the only significantly enriched terms were directly related to ‘translation’ or ‘ribosome assembly’ (hereon all simply referred to as ‘translation’ related – Fig. 2b and supplementary table S2a) that together comprised 28 of the differentially expressed proteins (Fig. 2c). Of these translation related proteins, 23 exhibited significantly decreased levels upon p38-MAPKi and 5 were upregulated (Fig. 2c). Hence, p38-MAPKi during early blastocyst maturation significantly dysregulates a specific cohort of translation associated proteins, mostly resulting in their reduced expression.

**Figure 2:**
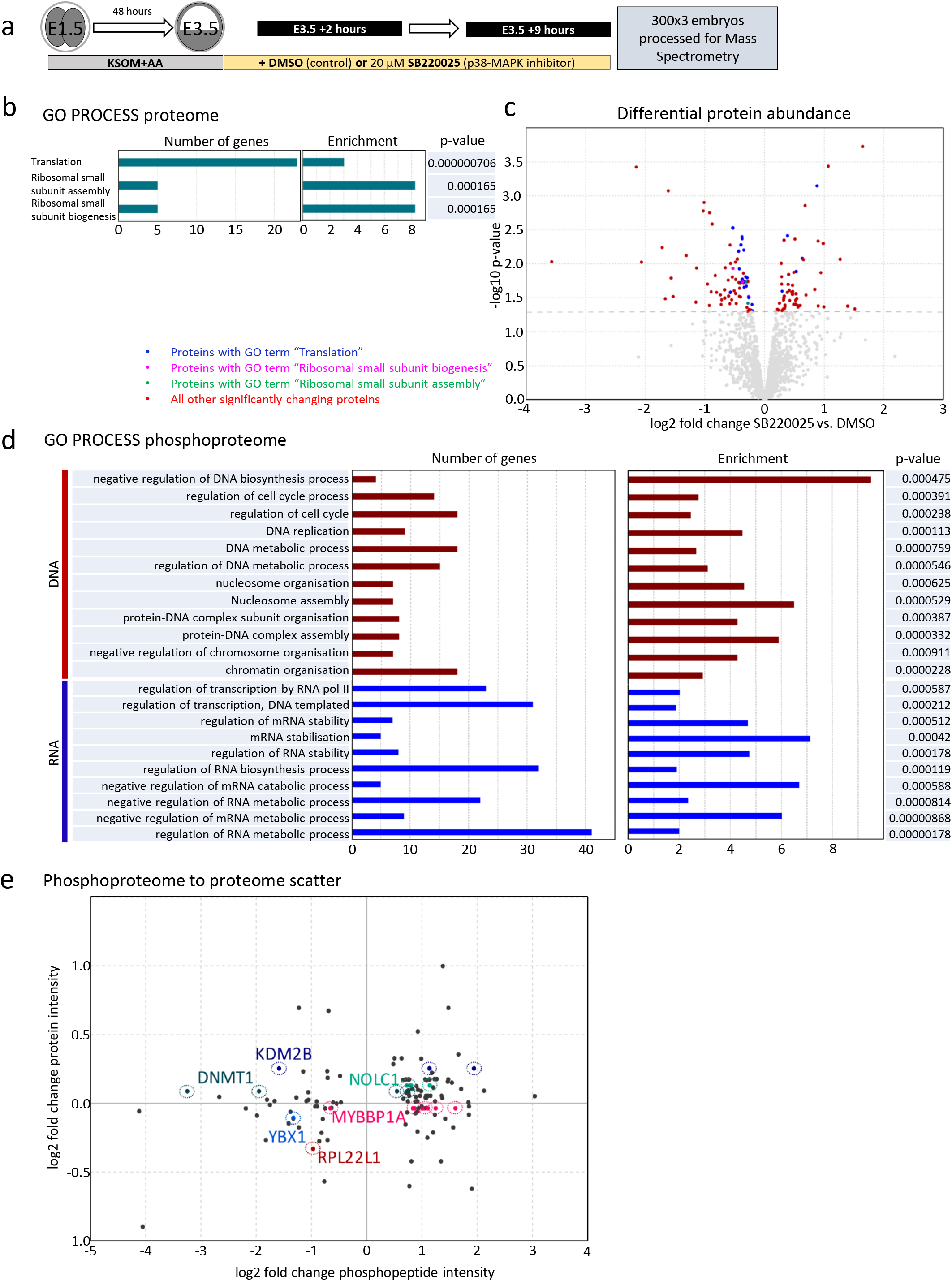
Proteomic and phosphoproteomic analyses of the effect of p38-MAPKi in developing blastocysts. a) Experimental design of sample collection of control (DMSO) and p38-MAPKi (SB220025) blastocysts after 7 hours chemical exposure (at E3.5 +9h) prior to mass spectrometric analysis of the (phospho)proteome (300 blastocysts per condition in biological triplicates). b) Most significantly enriched gene ontology terms of differentially detected proteins between control and p38-MAPKi blastocysts (full list of terms in supplementary table S2a). c) Volcano plot of the detected differential proteome expression dynamics associated with p38-MAPKi. Proteins belonging to the indicated significantly enriched GO terms are highlighted in blue (translation), purple (ribosomal small subunit biogenesis) and green (ribosomal small subunit assembly) – note there is overlap (see supplementary table S2a & b); the remainder in (red) with insignificant changing proteins in grey. d) Significantly enriched gene ontology terms for differentially detected phosphopeptides between control and p38-MAPKi blastocysts (full list of terms in supplementary table S2c). e) Phosphoproteome to proteome scatter plot for mass spectrometry detected peptides demonstrating <1.3-fold difference in abundance in the general proteome and >1.5-fold change in phosphorylation level. Candidates of interest (based on literature research) are highlighted (and were assayed for PrE phenotypes in clonal siRNA microinjection mediated loss of function assays – *e.g.* MYBBP1A, Fig. 4).

A similar interrogation of acquired phosphoproteome data revealed 156 significantly differentially enriched phosphopeptides between control and p38-MAPKi blastocysts (99 enhanced and 57 depleted); theoretically representing both direct p38-MAPK substrates and other indirect downstream effectors. Collectively, the differentially enriched phosphopeptides represented 110 individual proteins (supplementary table S2d). The detected changes in protein phosphorylation status corresponded to 59 proteins exhibiting solely enhanced phosphorylation, 41 proteins in which it was solely depleted and 10 proteins with both enhanced and depleted levels of phosphorylation at differing individual sites. We again performed an ontological analysis on genes identified in this phosphoproteomic screen and revealed an abundance of enriched terms related to RNA regulation and stability (Fig. 2d & supplementary table S2c) that resonated with those translation related terms identified from the interrogation of the general proteome (Fig. 2b), collectively suggesting p38-MAPKi may potentially impair protein synthesis during early blastocyst maturation. We also noted enriched terms related to cell cycle regulation, DNA replication and chromatin modification (Fig. 2d & supplementary table S2c) that is perhaps unsurprising given the understood wide range of p38-MAPK effectors^19^. To further refine the list of differentially phosphorylated proteins we removed any detected phosphopeptides theoretically mapping to proteins not detected in the general proteome. Next, we filtered out those detected sites that demonstrated less than 1.5 fold change in phosphorylation, retaining those with an equal to, or greater than, 1.5 fold change. From the remaining list, we retained those sites that exhibited a less than 1.3 fold change in their general abundance in the overall proteome (Fig. 2e & supplementary table S2d). We anticipated this would enrich for differentially expressed phosphoproteins with the highest impact on the observed phenotype, most closely associated with p38-MAPK regulation. Based on these criteria, we selected six target phosphoproteins that we functionally tested in clonal loss-of-function experiments (highlighted in Fig. 2 e, and expanded below). Hence, an assay of the p38-MAPKi sensitive phosphoproteome during the early blastocyst maturation period reveals specific alterations in the phosphorylation of multiple proteins, amongst which are an enriched population of metabolic RNA regulators.

In order to complement our (phospho)proteomic analysis we also conducted transcriptomic analyses of developing blastocysts under p38-MAPKi conditions. Accordingly, we processed mRNA from 30 embryos per condition at three different time-points during early blastocyst maturation; informed by our initial analyses of the p38-MAPKi sensitive window (Fig. 1)^25,26^. Consequently, the transcriptome was analysed after E3.5 +4h, +7h and +10h of p38-MAPKi and compared to that of control treatments (Fig. 3a). We detected 34 significantly differentially expressed gene transcripts at the +4h time-point, increasing to 1240 genes at the +7h time-point and reducing to 480 by the +10h time-point; only 10 differentially expressed genes overlapped between all three inhibition regimes (Fig. 3b & supplementary tables S3a (+4h), (+7h) and (+10h)). These data indicate profound changes in the p38-MAPK regulated transcriptome that are maximally centred around the E3.5 +7h time-point (Fig. 3b – supplementary table S3a (+7h)). Analysing the differentially expressed cohort of genes from this +7h time-point, we detected both upregulated and downregulated genes (supplementary table S3b (Clusters I, II and III)). When we performed hierarchical clustering of this cohort across all treatments and time-points (Fig. 3c), we identified three clusters of co-regulated genes (comprising >20 genes – note similar clustering of differentially regulated cohorts identified at E3.5 +4h and E3.5 +10h is provided in the supplementary information, Fig. S1 & tables S7). Cluster I represented genes whose transcript levels began to decrease after p38-MAPKi at the E3.5 +4h time-point, were maximally repressed at E3.5 +7h but returned to levels equivalent to control by E3.5 +10h. Gene transcripts in cluster III demonstrated a similar but inverted trend. Namely, at E3.5 +4h p38-MAPKi initiated an increase in transcript expression that was again maximal by E3.5 +7h and returned to control levels by E3.5 +10h. Cluster II was comprised of similarly enhanced transcripts levels but was distinguished from cluster III in that these did not return to the same basal control level by E3.5 +10h and remained elevated (supplementary table S3b (Cluster II and III)). Thus, p38-MAPKi was associated with the induction of large-scale transcriptome dynamics that peaked at the E3.5 +7h time-point and (with the exception of the enhanced gene transcripts represented in cluster II) were largely compensated back to the levels observed in control blastocysts by the E3.5 +10h time-point. However, notwithstanding such compensation, it is important to note p38-MAPKi throughout this same period, or simply in the identified minimal window of sensitivity, is still sufficient to impair PrE differentiation by E4.5 (Fig. 1 and^25,26^). Hence, the observed p38-MAPKi mediated transcript level changes may indeed be compensated by E3.5 +10h but the eventual PrE deficient phenotype persists, potentially due to impaired mRNA translation (see below). Interestingly, we mostly did not observe significant changes in the expression of these dynamically regulated genes between each of the three control conditions (*i.e.* +DMSO at E3.5+ 4h, +7h and +10h), indicating such gene mRNA levels are not normally subject to regulation during this period. Combined, these data reveal a profound, yet largely transient, sensitivity of specific cohorts of gene mRNA expression to the activity of p38-MAPK during the early blastocyst maturation developmental window. Remarkably, when we searched for gene ontological terms associated with this described transcriptomic dynamism (Fig. 3d & supplementary tables S3c), we again identified translation related terms as most significantly enriched (mostly reflecting upregulated expression – clusters III, Fig. 3c); followed by mitochondrial respiration (also upregulated – cluster III, Fig. 3c) and splicing linked terms (representing both upregulated and downregulated transcripts – clusters I and III, respectively, Fig. 3c). From the 82 differentially expressed translation related genes identified in the E3.5 +7h time-point (supplementary table S3d), we found 79 were upregulated by p38-MAPKi (supplementary tables S3d). Interestingly, 13 of those 79 upregulated gene transcripts (mainly representing small ribosomal subunit proteins) had also been identified as significantly down-regulated in the general proteome (Fig. 3e & supplementary tables S3e). Thus, these data suggest the accumulation of translation related mRNAs caused by p38-MAPKi (maximal at E3.5 +7h) might be related to impaired ribosomal assembly and/or functional protein translation. Moreover, they also indicate a potential positive feedback mechanism, culminating in the transiently enhanced levels of translation/ribosome related gene transcripts observed (Fig. 3c & d).

**Figure 3:**
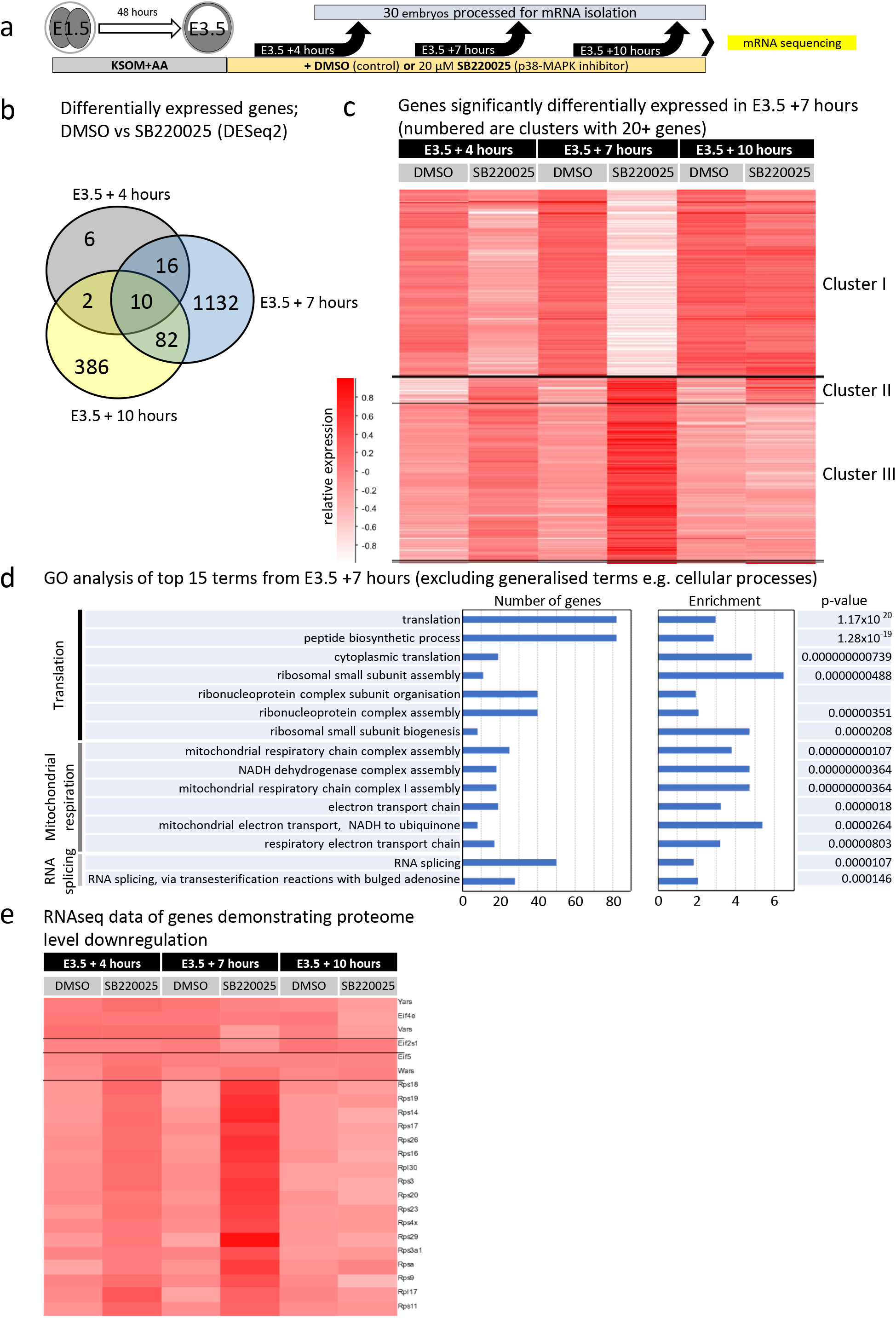
Temporal transcriptomic analysis of the effect of p38-MAPKi during early blastocyst maturation. a) Experimental design for transcriptome sample collection following control (DMSO) or p38-MAPKi (SB220025) from E3.5 for +4h, +7h or +10h (biological duplicates of 30 blastocysts per condition per time-point employed). b) Venn diagram representing number of differentially expressed gene mRNAs (DESeq2 analysis) and their overlap between control and p38-MAPKi blastocysts in the three selected time-points (*i.e.* E3.5 +4h, +7h and +10h) (gene lists provided in supplementary tables S3a). c) Hierarchical clustering heat-map depicting the expression of significantly changing gene mRNAs elicited by p38-MAPKi at the +7h time-point and the status of the transcript levels of those same genes at the +4 and +10h time-points; forming three distinct expression clusters. (gene lists, per cluster, provided in supplementary tables 3b). d) Statistically enriched gene ontology analysis of the top 15 terms identified in the p38-MAPK regulated transcriptome at the +7h time-point (excluding generalised terms – full list of terms in supplementary table S3c). e) Hierarchical clustering heat-map of gene mRNAs, originally identified as downregulated at the protein level and associated with translation-related gene ontology after p38-MAPKi (Fig. 2b & c), at the three assayed ±p38-MAPKi early blastocyst time-points (supplementary table S3e).

Overall, the combined (phospho)proteomic and transcriptomic analyses point to regulation of translation as a primary function of p38-MAPK activity during early blastocyst maturation. Moreover, when considered in the wider context, such regulation potentially creates a permissive state for subsequent ICM cell fate derivation (particularly PrE differentiation), that we focus upon in the remainder of this study.

### MYBBP1a has a prominent role in preimplantation embryonic development

As referenced above, we applied extra significance filters to the results of our phosphoproteomic screen in an effort to identify phosphoproteins either directly regulated by p38-MAPK or functionally related post-translational control (Fig. 2e). We further used literature precedent to identify genes previously shown or hypothesized to have roles in developmental pluripotency and/or differentiation, with special emphasis on translation. Thus, we selected a total of six candidate genes for further loss-of-function studies in the preimplantation mouse embryo; *Dnmt1, Kdm2B, Ybx1, Mybbp11, Rpl22l1 and Nolc1* (Fig. 2e). Accordingly, we employed specific siRNA constructs (or non-targeting controls) against the candidate transcripts to elicit gene expression knockdown in marked preimplantation embryo cell clones. This was achieved by co-microinjection of appropriate siRNAs with recombinant *in vitro* transcribed mRNA encoding a histone H2B-RFP fusion reporter protein in one cell of 2-cell stage (E1.5) embryos, resulting in a fluorescently marked clone of cells comprising half the embryo. Embryos were then cultured until the late blastocyst (E4.5) stage and immuno-fluorescently stained for the ICM cell fate markers NANOG (EPI) and GATA4 (PrE). Late blastocysts derived from embryos microinjected in both blastomeres at the 2-cell stage were used as a source of mRNA for qRT-PCR analyses to assess the extent of specific candidate transcript knockdown at E4.5 (Fig. 4a).

**Figure 4:**
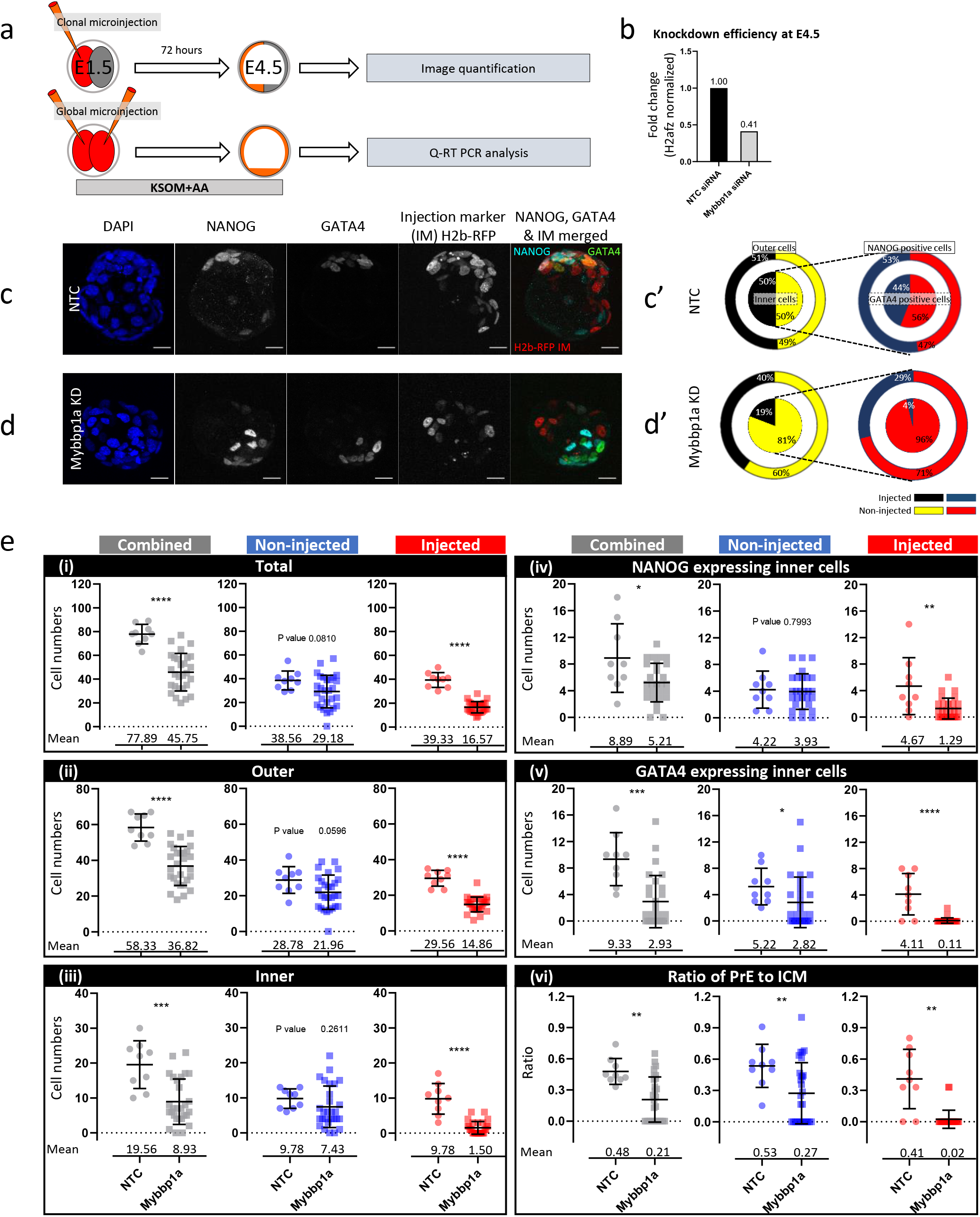
Analysis of the role of MYBBP1A (candidate target of p38-MAP kinase activity) on preimplantation embryonic development and ICM cell fate specification. a) Experimental design to determine the efficiency of siRNA mediated *Mybbp1a* gene mRNA knockdown in microinjected embryos cultured to the equivalent late blastocyst (E4.5) stage (lower panel) and to assay the contribution of marked *Mybbp1a* knockdown clones to late blastocyst cell lineages (upper panel). b) qRT-PCR derived relative *Mybbp1a* transcript levels (normalised to *H2afz* mRNA levels) between embryos injected with non-targeting control (NTC) siRNA and siRNA specific for *Mybbp1a* mRNA. c) Confocal micrograph z-projections of an exemplar late (E4.5) stage blastocysts initially microinjected (in one blastomere at the 2-cell stage) with NTC siRNA (n=9) plus recombinant *H2b-RFP* fluorescent reporter mRNA (identifying the clonal progeny of the injected cell). Individual DAPI (blue pan-nuclear stain; total number of cells), NANOG (greyscale; EPI cells) and GATA4 (greyscale; PrE cells) channel micrographs, plus a merged NANOG (cyan), GATA4 (green) and H2B-RFP (microinjected clone) image are show (scale bar = 20μm). (c’) Target diagrams describing average percentage contribution of NTC siRNA microinjected and nonmicroinjected clones to either outer and inner cell populations (black and yellow targets) or mutually exclusive NANOG or GATA4 expressing ICM cell lineages (red and blue targets) d) & d’) As in (c) and (c’) but after microinjection of *Mybbpa1* specific siRNA (n=28). e) Scatter plots describing the cell number contribution from non-microinjected (blue) and microinjected (red) cell clones (plus the combined number – grey) of *NTC/Mybbp1a* siRNA treated embryos to: (i) total cell count, (ii) outer cells, (iii) inner cells), (iv) NANOG only expressing inner cells, (v) GATA4 only expressing inner cells, plus (vi) the ratio of PrE (GATA4 positive) cells to overall ICM size. Means and standard deviation highlighted. Collated cell count data for each embryo used in each scatter plot is provided in supplementary table S4.

Of the six short-listed candidates, it was the targeting of *Mybbp1a* that revealed the most substantial developmental impact, despite the relatively modest transcript knockdown efficiency of 59% (Fig. 4b); the effects of other candidate gene knockdowns are summarised in the supplementary information (table S8). MYBBP1A is a predominantly nucleolar protein with reported roles in the negative regulation of RNA polymerase I dependent rRNA gene transcription, co-transcriptional rRNA processing and ribosome biogenesis^33,34^, p53 tetramerization, cell senescence and apoptosis^35,36^, plus negative regulation of *Hoxb2* gene transcription^37^. Additionally, the crossing of heterozygous *Mybbp1a* null mice is unable to support the development of *Mybbp1a* homozygous null blastocysts^38^; implying an important, yet non-defined, role in preimplantation development. Our clonal analysis of fluorescently marked *Mybbp1a* knockdown cells revealed a 19% contribution to late blastocyst (E4.5) stage ICM and 40% to outer TE cells; compared to 50% and 51%, respectively in control siRNA injected embryos (Fig. 4c-e). Although the reduced ICM/outer cell contribution of the *Mybbp1a* knockdown clone was marked by an overall reduction in its total cell number size when compared to the equivalent microinjected clones of control siRNA treated embryos (comprising an average of 16.57, standard deviation (SD) 6.20, versus 39.33, SD 4.71, cells, respectively) or its non-injected sister clone (Fig. 4e). However, despite the smaller sized *Mybbp1a* knockdown clone population exhibiting reduced contribution to the ICM, those inner allocated cells rarely, if at all, contributed to GATA4+ PrE (only representing an average of 0.11 cells, SD 0.42, per embryo, or 4% of the total PrE population). Conversely, such clones readily contributed to NANOG+ EPI (comprising an average of 1.29 cells, (SD 1.56), per embryo or 29% of the total EPI cell number). Contrastingly, in control siRNA treated embryos the contribution of both microinjected and non-microinjected clones between both ICM and outer cells, plus the EPI and PrE lineages, was always observed to be statistically equal (Fig. 4 c-e); indicating a lack of ICM clonal bias potentially introduced by the microinjection procedure itself. Interestingly, the clonal *Mybbp1a* knockdown also mildly affected the non-microinjected clone, leading to small yet statistically insignificant reductions in total, outer and inner cell number. The number of GATA4 positive PrE cells within the non-microinjected clone was, however, significantly reduced (averaging 2.82, SD 3.83, versus 5.22, SD 2.77, cells in the equivalent clone of control siRNA treated embryos) but the number of EPI cells was statistically unaffected (averaging 3.93, SD 2.67, versus 4.22, SD 2.77, cells). Thus, suggesting that the overall reduced cell number of the *Mybbp1a* clonal knockdown in blastocyst may also negatively affect PrE differentiation in the non-injected clone, *per se* (Fig. 4 c-e & supplementary tables S4).

In summary, siRNA mediated clonal knockdown of *Mybbp1a,* a known regulator of rRNA transcription and processing, and identified in this study as a potential p38-MAPK substrate during early blastocyst maturation (Fig. 2e), biases ICM resident clones against PrE differentiation. Hence, implicating MYBBP1A as an ICM cell fate regulator acting downstream of required p38-MAPK activity.

### p38-MAPK inhibition disrupts rRNA processing and reduces active translation and transcription during early blastocyst maturation

The above described (phospho)proteomic and transcriptomic screens, plus the functional validation of *Mybbp1a*, a reported player in translation regulation mechanisms^33^, strongly supported the hypothesis that p38-MAPK regulated translation is a major contributor to germane blastocyst maturation. Thus, we wanted to directly assay the effect of p38-MAPKi on translation. Accordingly, we utilised the O-propargyl-puromycin (OPP) incorporation assay (that permits fluorescent labelling of nascent translated polypeptides, that can be quantified after confocal microscopy imaging) to measure blastocyst translation during a 10 hour period of early blastocyst maturation (starting from E3.5), ± p38-MAPKi (Fig. 5a). As was expected, we recorded a near 50% decrease in global protein synthesis upon p38-MAPKi (Fig. 5b).

**Figure 5:**
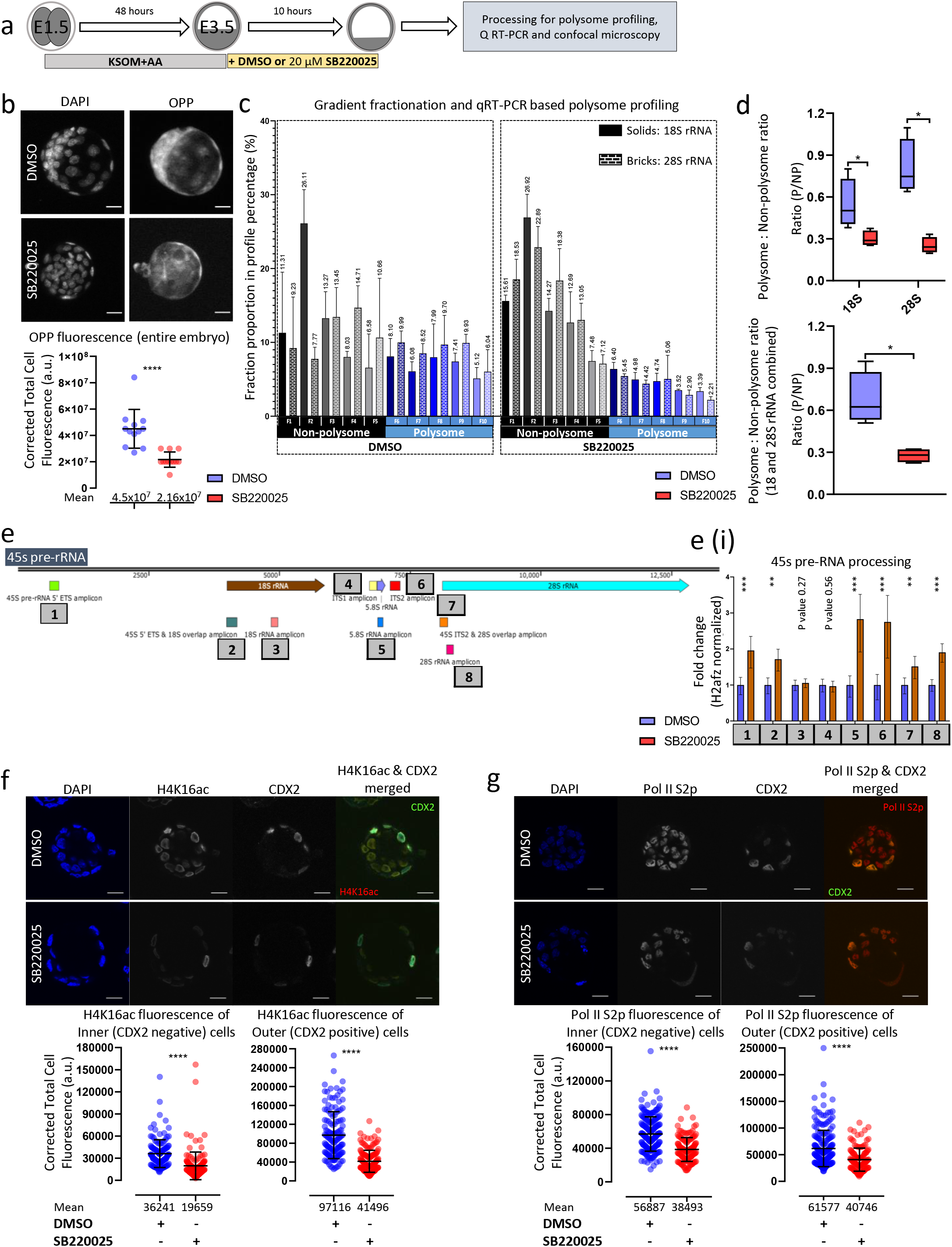
The effect of p38-MAPKi on active translation, rRNA processing and transcriptional landscape in developing blastocysts. a) Experimental design for analysing the role of p38-MAPK activity on developmental translation and transcription in maturing blastocysts (between E3.5-E3.5 +10h). b) Exemplar confocal z-projection micrographs (upper) of E3.5 +10h blastocysts in control (DMSO; n=12) and p38-MAPKi (SB220025; n=12) conditions, depicting cell nuclei staining (DAPI) and *de novo* translation (after OPP incorporation); scale bar = 20μm. Scatter plot quantification of Corrected Total Cell Fluorescence (CTCF) of OPP incorporation between the two conditions (lower); means and standard deviation highlighted (supplementary table S5a). c) qRT-PCR analyses quantifying 18S and 28S rRNA levels in each of ten Scarce Sample Polysome (SSP) profiling fractions, representative of polysome associated (F6-F10) and non-polysome associated (F1-F5) rRNA transcripts, obtained from both control and p38-MAPKi blastocysts (E3.5 +10h). Individual bars represent the proportion to the overall level of detected rRNA transcripts attributable to a given fraction. Data generated from four sets of 10 embryos each, per condition; error bars denote standard deviations. d) Ratio of polysome to non-polysome quantified rRNAs (as derived from data in Fig. 5c) for 18S and 28S separately (top panel) and combined (bottom panel). Data presented as interleaved box and whiskers plot, with whiskers depicting minimum and maximum values and box plot including medians and the 25^th^ and 75^th^ percentiles. e) Schematic representation of the 45S pre-rRNA, highlighting the qRT-PCR amplicons assayed and quantified in order to analyse rRNA processing status ±p38-MAPKi (E3.5 +10h). i. Bar graphs representing mean qRT-PCR quantified fold changes of each specific amplicon (as normalised against *H2afz* cDNA) in p38-MAPKi (red) versus control blastocysts (DMSO – blue), plus standard deviations. f) Exemplar single z-stack micrographs (upper) of E3.5 +10h blastocysts after control (DMSO; n=13) and p38-MAPKi (SB220025; n=15) treatment. Individual DAPI (blue pan-nuclear stain; total number of cells), H4K16ac (greyscale; post-translational histone chromatin mark associated with transcriptional activity) and CDX2 (greyscale; marking outer TE cells) channel micrographs, plus a merged H4K16ac (red) and CDX2 (green) image are show (scale bar = 20μm). Scatter plots (lower) quantifying per cell nuclei H4K16ac (CTCF) levels in control (DMSO) and p38-MAPKi (SB220025) blastocysts, differentiated between inner (n=129 for DMSO and n=143 for SB220025) and outer (n=136 for DMSO and n=145 for SB220025) cell populations; means and standard deviations depicted (supplementary table S5b). g) As in (f) but describing expression levels of RNA polymerase II C-terminal domain phosphoserine 2 (pol II S2p) protein. Inner cell (n=187 for DMSO and n=135 for SB220025) and outer cell (n=209 for DMSO and n=123 for SB220025) quantification, with mean and standard deviation shown (supplementary table S5c).

We next sought to verify the observed reduction in active translation using a polysome profiling approach. Therefore, we adopted our recently reported protocol of Scarce-Sample Polysome (SSP) Profiling^39^, specifically optimised for samples of limiting size, as a readout of active translation in early blastocysts ± p38-MAPKi. We procured ten fractions representing individual ribosomal subunits, monosomes and polysomes, from 10 blastocysts per condition in quadruplicate. The fractions were then processed for total RNA extraction, cDNA synthesis and analysed by qRT-PCR to specifically assay 18S and 28S rRNA levels. We observed a discernible shift in the distribution of both 18S and 28S rRNA abundance towards SSP fractions indicative of free rRNAs or association with individual ribosomal subunits/monosomes (fractions 1-5) versus denser fractions indicative of polysomes (fractions 6-10) in the p38-MAPKi treated embryos (Fig. 5c). These combined rRNA enrichment/fractionation data can be expressed as calculated polysome to non-polysome ratios, to provide a further indication of the state of active blastocyst translation (*i.e.* higher ratios indicative of enhanced active translation). As can be seen, we observed a significant decrease in the ratio of polysomes in p38-MAPKi blastocysts (Fig. 5d bottom panel) that was greater in magnitude for 28S compared to 18S rRNA targeted assays (Fig. 5d top panel). Therefore, we conclude these and the OPP-derived data as confirming reduced active translation in mouse blastocysts under p38-MAPKi during the early stages of maturation.

It is reported that MYBPP1A interacts with HTATSF1, a factor known to be involved in transcriptional and post-transcriptional regulation/processing of rRNA and mRNA. HTATSF is also a confirmed regulator of pluripotency and has a role in the transition between pre- to post-implantation embryonic development via the regulation of protein synthesis^40^. Thus, using qRT-PCR, we assayed the effect of p38-MAPKi on 45S pre-rRNA processing, during the same 10 hour early blastocyst developmental window described above (note, we used the same primer sequences as described in the HTATSF1 report of Corsini *et al.,* to assay specific regions of the 45S pre-RNA indicative of the extent of processing^40^). This analysis revealed p38-MAPK does regulate rRNA processing during early blastocyst maturation (Fig. 5e & e (i)). Specifically, we observed enhanced levels of amplicons indicating impaired processing. For example, the internal transcribed spacer ITS2 region, normally removed from the 45S pre-rRNA during processing, was detected at levels nearly 3-folds higher than in control blastocysts. Similarly, levels of the 5’ external transcribed spacer (ETS) were also significantly increased. Moreover, regions representative of the overlap between ITS2 and the mature 28S rRNA coding sequence, plus the overlapping region between the ETS and 18S rRNA, were also significantly increased; indicating impaired 45S pre-rRNA processing under p38-MAPKi conditions. Using primers specific to regions of the 45S pre-rRNA that correspond to the mature (appropriately processed) rRNA transcripts, we observed significant increases in both the 28S and 5.8S rRNAs, that presumably reflect the increases in unprocessed 45S pre-rRNA intermediates. However, a similar assay of the 18S rRNA transcript was not associated with an increase compared to control, suggesting steady state levels of the mature and processed 18S rRNA were not significantly affected. This was also reflected in the fact the levels of ITS1 also remained unchanged and distinguishes such p38-MAPKi data from studies of *Htatsf1* genetic knockouts in which ITS levels increase (and ETS levels fall)^40^. Notwithstanding such differences, the data presented here confirm p38-MAPKi during early blastocyst maturation is associated with the accumulation of 45S pre-RNA processing intermediates, which is concomitant with a reduction in polysome formation (potentially via impaired ribosome biogenesis) and reduced active translation.

In the context of peri-implantation embryonic development and *in vitro* pluripotency, recent reports have drawn important correlative links between the general translational status and permissive transcriptional landscapes^41,42^. Therefore, we asked if the observed p38-MAPKi mediated defects in protein synthesis were also associated with negatively affected transcription (Fig. 5f & g). Consistently, we observed a significant reduction in the expression levels of histone H4-lysine 16 acetylation (H4K16ac, a post-translational mark of actively transcribed chromatin^43^). The reduction was revealed in both inner and outer cells, as demarcated by the expression of CDX2 (an outer cell TE marker^44^). Similarly, immuno-fluorescent staining for actively elongating RNA polymerase II (using an antibody that recognises RNA polymerase II phosphorylated on serine 2 of the C-terminal domain – RNA pol II S2p^45^) also showed a marked decrease upon a 10 hour period of p38-MAPKi from the E3.5 stage.

Hence, p38-MAPKi during early blastocyst maturation, known to negatively affect subsequent ICM cell fate and specifically PrE differentiation (described above^25,26^), is associated with impaired translation, polysome formation, rRNA processing and a less permissive transcriptional landscape.

### p38-MAPK and mTOR, a key regulator of protein translation, have partially overlapping roles in regulating preimplantation embryonic development

Although p38-MAPK activity has been reported to regulate protein synthesis in other contexts^46^, mTOR mediated translation regulation represents the most understood and extensively studied pathway, particularly in development^47^. Observations from studies of cancer biology indicate a possible convergence of both pathways towards positive regulation of eIF4E-sensitve cap-dependent mRNA translation^48,49^. Therefore, the generally significant role mTOR plays in translation regulation, coupled to a recent report describing blastocyst diapause caused by attenuated translation and transcription after mTOR inhibition^41^, prompted us to investigate our observed p38-MAPK phenotypes in conjunction with mTOR activity. We first sought to directly compare p38-MAPKi phenotypes with those elicited by TORIN1 mediated pharmacological inhibition of mTOR activity^50^. Early blastocysts (E3.5) were cultured in the presence of control DMSO, TORIN1 (inhibiting mTOR) and SB220025 (inhibiting p38-MAPK) for 24 hours (to the late blastocyst/E4.5 stage) and immuno-fluorescently stained for ICM cell lineage markers (NANOG – EPI and GATA4 & GATA6 – PrE; Fig. 6a). We found that mTOR inhibition, like p38-MAPKi significantly increased the proportion of NANOG and GATA6 co-expressing cells within the ICM (Fig. 6b-d & p), a hallmark of unspecified ICM^10^ and potentially that of a diapaused state. Likewise, both treatment groups comprised blastocysts with significantly fewer PrE cells *(i.e.* defined as cells solely expressing GATA4 – Fig. 6b-d, n & o), confirming both our previous p38-MAPKi observations^25,26^ and the reported diapaused state of mTOR inhibited blastocysts^41^. However, despite these similarities, we observed mTOR inhibited blastocysts had significantly fewer EPI cells (and overall ICM cells) than counterpart p38-MAPKi and control embryos (*i.e.* defined as cells solely expressing NANOG – Fig. 6b-d, l & m). This indicates that mTOR inhibited blastocysts are indeed diapaused and reflect the early blastocyst developmental stage, characterised by an unspecified ICM (something not confirmed in earlier studies), at which TORIN1 was first administered. Conversely, the fact that p38-MAPKi blastocysts were able to specify EPI cells yet retain a population of unspecified ICM cells and were impaired in PrE formation, indicates the effect of p38-MAPKi (and the associated uncovered translational defects) is centred on PrE differentiation. Therefore, it is unlikely p38-MAPKi is a simple phenocopy of the developmental diapause induced by mTOR inhibition. This conclusion is supported by the fact that continued blastocyst culture beyond E4.5 under p38-MAPKi conditions results in embryo death by E5.5-E7.5 but continued mTOR inhibition beyond this point is still associated with intact embryos (see supplementary Fig. S2), indicative of true diapause^41^. Therefore, whilst blastocyst maturation phenotypes associated with mTOR and p38-MAPKi share similarities they are not complete phenocopies. Although this does not rule out the possibility that p38-MAPK and mTOR functionally converge/cooperate in the limited context of PrE differentiation.

**Figure 6:**
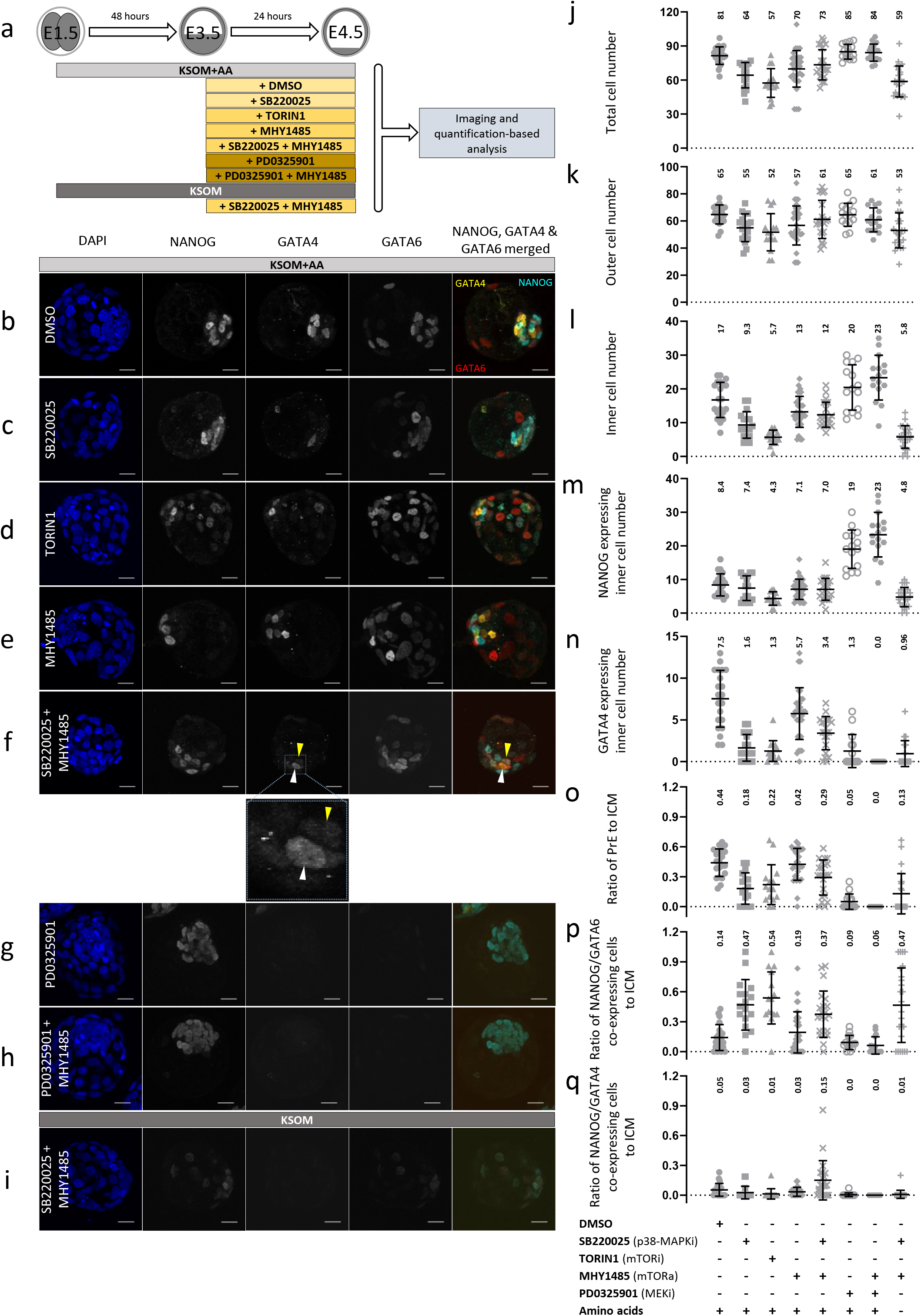
Functional interplay between mTOR and p38-MAPK during blastocyst maturation and PrE differentiation. a) Experimental schematic for analysing potential functional overlap of mTOR and p38-MAPK on PrE differentiation (assaying ICM lineage marker protein expression) during blastocyst maturation; whereby E3.5 stage blastocysts were cultured for 24 hours in media (± amino acids/AA) containing the indicated pharmacological supplements (vehicle control – DMSO; p38-MAPK inhibitor – SB220025; mTOR inhibitor – TORIN1; mTOR activator – MHY1485; MEK1/2 inhibitor – PD0325901) or combinations thereof. b-i) Exemplar confocal z-series projections of blastocysts cultured under the stated conditions described in (a) and stained for DAPI (blue) NANOG, GATA4 and GATA6 protein expression (cyan, yellow and red in provided merged image); scale bar = 20μm. (b) DMSO (n=21), (c) p38-MAPKi, SB220025 (n=17), (d) mTOR inhibition, TORIN1 (n=15), (e) mTOR activation, MHY1485 (n=32), (f) p38-MAPKi and mTOR activation, SB220025 + MHY1485 (n=22), (g) MEK inhibition, PD0325901 (n=15), (h) MEK inhibition and mTOR activation, PD0325901 + MHY1485 (n=16) and (i) p38-MAPKi and mTOR activation, SB220025 + MHY1485 in KSOM non-supplemented with amino acids (n=24). Note in panel (f), and the expanded view, white arrowheads highlight differentiated PrE cells expressing GATA4 (and GATA6) but not NANOG, whereas yellow arrowheads show ICM cell co-expressing GATA4 and NANOG (that are comparatively enriched in this condition). j)-q) Scatter plots of total, outer and inner blastocyst cell number (evidenced by DAPI staining) and the numbers of NANOG, GATA4 and GATA6 expressing ICM cells under stated culture conditions described in (a) (with mean and standard deviation highlighted). *i.e.* j) total cells, k) outer cells, l) inner cells, m) NANOG expressing inner cells and n) GATA4 expressing inner cells; plus o) ratio of GATA4 expressing, NANOG non-expressing, PrE to total ICM cells, p) ratio of NANOG and GATA6 co-expressing cells to total ICM cells and q) ratio of NANOG and GATA4 co-expressing cells to total ICM cells. Collated cell count data for each embryo used in each scatter plot is provided in supplementary table S6a and results of statistical tests in table S6b.

Therefore, we next asked if p38-MAPKi mediated PrE specific differentiation phenotypes could be ameliorated by pharmacological activation of mTOR. Accordingly, we utilised the potent mTOR activator MHY1485^51^, that when provided in isolation to maturing blastocysts (E3.5-E4.5) has no discernible effect on development and ICM cell fate derivation, compared to control embryos (Fig. 6a, e & j-q). However, when mTOR activator was administered in combination with p38-MAPKi, the number of GATA4 expressing PrE cells was significantly increased compared to the number observed after p38-MAPKi alone, although not reaching the levels seen in control blastocyst (Fig. 6a, b, f & n). This increase was also reflected in a reduced contribution of unspecified ICM cells, co-expressing NANOG and GATA6 (Fig. 6p). We also noted an accompanying increase in total embryo cell number, which resulted from modest increases in the number of outer and inner cells, although there was no change in the number of NANOG alone expressing EPI inner cells (also reflected in the PrE to ICM ratios – Fig. 6j-m). Interestingly, we also observed a significant population of ICM cells that co-expressed NANOG and GATA4, that in all other conditions were seldom observed (Fig. 6f & q). This suggests that the p38-MAPKi mediated defect in PrE differentiation, that normally results in a unique population of unspecified PrE progenitors, can be partially overcome by activation of mTOR but without an accompanying decrease in NANOG expression. In conclusion, blastocyst maturation PrE deficient phenotypes caused by p38-MAPKi can be partially rescued by concomitant activation of mTOR.

The FGF4-FGFR-MEK-ERK cascade is considered the quintessential ICM cell fate determining signalling mechanism in mouse blastocysts^52^. We have previously reported that PrE deficient phenotypes caused by sole FGF-receptor (FGFR) inhibition can be rescued by activation of endogenous p38-MAPK^25^. However, informed by our latest observations, we assayed if PrE differentiation phenotypes induced by MEK inhibition (using PD0325901^53^), presumably functioning downstream of activated FGFR, were also sensitive to mTOR activation. Whilst pharmacological inhibition of MEK/ERK (E3.5-E4.5) resulted in the well-defined phenotype of largely pan-ICM expression of NANOG alone, there was no PrE rescue effect caused by the additional activation of mTOR; indeed the effect of MEK inhibition was augmented (Fig. 6g, h & j-q). We therefore, conclude that the role p38-MAPK plays in ICM cell fate specification, via translation regulation, is functionally and mechanistically distinct from the role played by the MEK/ERK pathway.

We previously reported p38-MAPK also acts to counter amino acid depletion induced oxidative stress in the developing mouse blastocyst; as revealed by significantly more severe ICM cell fate phenotypes, affecting both EPI and PrE derivation, when p38-MAPKi was combined with a lack of exogenous amino acid media supplementation^26^. Thus, we speculated if activating mTOR could also alleviate such enhanced effects of p38-MAPKi associated with amino acid depletion (E3.5-E4.5). Unlike in the presence of supplemented amino acids, we found blastocysts cultured under such conditions did not demonstrate any induced GATA4 inner cell expression or increased inner cell number; nor was there any increase in NANOG alone expressing EPI cells (Fig. 6i & j-q). These data confirm the previously identified role of p38-MAPK in counteracting oxidative stress is distinct from that described here in relation to mTOR activity and translation.

In conclusion, we have identified a novel developmental role for p38-MAPK in regulating translational output during the early stages of preimplantation mouse embryo blastocyst maturation. This role is required to allow a permissive environment for PrE differentiation and is at least partially integrated with the mTOR pathway.

## Discussion

The p38-MAPKs have well defined roles as stress activated kinases involved in inflammatory diseases, including cancers, and are also known to play developmental roles in species as evolutionarily distant as sea urchins, flies, frogs, zebrafish and mouse^20^. In preimplantation mouse embryonic development, p38-MAPK signalling was first defined to be necessary for blastocyst formation (*i.e.* progression past the morula stage)^22,23^. Genetic knockout studies have established that loss of the *Mapk14* gene has an embryonic lethal phenotype associated with defective extraembryonic placental tissue formation and insufficient nutrient transfer, with homozygous null mutant conceptuses being resorbed around E12.5^21^. Our own previous studies (also confirmed here) have demonstrated the importance of p38-MAPK activity in the derivation of the primitive endoderm^25,26^, itself the precursor of the supportive extraembryonic parietal and visceral endodermal membranes^54^. In this report, we have not only experimentally recapitulated our original p38-MAPKi mediated PrE differentiation/deficient phenotype (Fig. 1a(ii)) but have also confirmed a blastocyst expansion defect (Fig. 1b) characterised by initially unaffected expansion (first 3 hours), leading into a phase of stifled (hours 3-10) and then arrested expansion (Fig. 1c). The fact p38-MAPKi blastocysts (E3.5-E4.5) exhibit cavity expansion defects and have impaired PrE derivation (Fig. 1)^25,26^ resonates with a recent report describing a positive correlation between cavity expansion and EPI and PrE cell fate specification and sorting^12^. In their study, Ryan and colleagues showed the chemical modulation of blastocyst cavity size (using Ouabain – an inhibitor of the ATP1 channel responsible for cavity fluid accumulation^55–57^) during early blastocyst maturation (E3.5-E4.0), reduced the expression levels of both EPI (SOX2) and PrE (GATA4) marker proteins (assayed at E4.0). Additionally, they demonstrated mechanical deflation of the cavity during the same period, to maintain an approximately constant E3.5-like volume, impaired GATA4 expression but not SOX2 levels (at E4.0). Whilst the extent of cavity expansion impairment was more pronounced using the mechanical approach, both manipulation strategies also resulted in reduced sorting of the EPI and PrE progenitor populations, compared with controls/simulations^12^. It is striking that the E4.5 PrE differentiation defects associated with p38-MAPKi we report (here and previously^25,26^) closely resemble those in mechanically deflated blastocyst, although these were assayed at E4.0^12^; *i.e.* representing significantly fewer unsorted GATA4 positive ICM cells and overall reduced GATA4 expression, that in both cases is accompanied by overtly normal specification of the EPI (marked by robust expression of either NANOG or SOX2 alone). Our own observations of p38-MAPKi late blastocysts demonstrate the comparative lack of GATA4 positive ICM cells is accompanied by a compensatory population of unspecified progenitors co-expressing NANOG and GATA6, indicating a PrE specific differentiation defect^25,26^. It is tempting to speculate the existence of a similar population of NANOG and GATA6 co-expressing unspecified PrE progenitors in mechanically deflated blastocysts, although this was not directly assayed^12^. Thus, it is possible that the impaired blastocyst PrE differentiation phenotypes we observe upon p38-MAPKi are, at least in part, mediated by insufficient cavity expansion. In support of this hypothesis, we calculated the average cavity volume of p38-MAPKi blastocyst (cultured in the presence of the inhibitor from E3.5-E4.5) as 162.9pL, equivalent to an equatorial area of 3316μm^2^ (median value 111.9pL), which is similar to the average E3.5 stage (149pL) and mechanically deflated E4.0 blastocyst (78pL) values reported^12^. Therefore, we can conclude that p38-MAPK activity is required for appropriate blastocyst volume expansion and that this has the potential for knock on consequences for PrE specification and differentiation.

The PrE specific p38-MAPKi mediated phenotype we describe could therefore potentially be conceptualised, in part, as a TE defect that is manifest in impaired blastocyst cavity expansion. Indeed mouse blastocyst inhibition of p38-MAPK activity has been reported to increase TE tight junction permeability and reduce *Aqp3* expression^23^, that together with the action of Na^+^/K^+^ pumps (such as ATP1) are required to initiate and maintain blastocyst cavity expansion and luminal pressure^23,58^ (that between E4.0 and E4.5 is reported to more than double^58^). Interestingly, our (phospho)proteomic and transcriptomic data indicate p38-MAPKi mediated downregulation of *Tjp1* at both the protein and RNA levels (~50% – supplementary data tables S2b and S3a (+7h)). ATP1 is comprised of ATP α1 and ATP β1 subunits and is acknowledged as the primary driver of cavity fluid filling^12,57^. Deletion of the α1-subunit gene (*Atp1a1*) does not prevent cavitation but it does cause peri-implantation lethality and is accompanied by cavity collapse after *in vitro* culture^59^. Whereas, siRNA mediated preimplantation embryo knockdown of *Atp1b1* expression prevents initiation of cavitation^57^. Contrary to previous reports^23^, our transcriptomic and proteomic analysis demonstrate a near 40% decrease in both mRNA and protein expression levels of *Atp1b1* after p38-MAPKi during the early blastocyst maturation stage, but we also confirm *Atp1a1* expression remains unchanged (supplementary data tables S2b, S3a (+7h) and S9). Taken in collective context, the impaired PrE differentiation caused by inhibition of p38-MAPK during blastocyst maturation does correlate with reduced cavity expansion, potentially caused by reductions in the expression of functionally related genes in the TE (*e.g. Tjp1* and *Atp1b1*). Whether this reflects targeted regulation of such specific genes or is related to the more general p38-MAPK mediated control of overall translation, identified and described in this report, is not clear. However, it is also important to note that PrE differentiation can occur in the absence of a TE/blastocyst cavity. For example, ICMs immunosurgically isolated from E3.5 stage mouse blastocysts are able to support formation of an outer layer of PrE within 24 hours (mirroring PrE differentiation in intact blastocysts)^60^ as can mouse ES cell derived ICM organoids (following a pulsed induction of *Gata6* expression)^61^.

Although, p38-MAPK regulation of blastocyst cavity size has the potential to influence the resolution of ICM cell fate (specifically PrE differentiation), our data do not necessarily rule out the existence of other contributory/parallel mechanisms that may act at the level of ICM cells themselves. It is well established that FGF4 based signalling from emerging EPI cells drives PrE differentiation from a decreasing pool of unspecified cells. Moreover, that this process is incremental and ensures a stable EPI:PrE cell number ratio throughout and requires activation of the mitogen activated kinase kinases MEK1/2 (MEK)^7,9^. We and others have comprehensively shown that pharmacological inhibition of MEK (either alone or in combination with FGFR inhibition) throughout the entire blastocyst maturation period completely blocks PrE differentiation (resulting in pan-ICM expression of EPI markers). Furthermore, that removal of MEK inhibition during this period at any point prior to ~E4.25 is permissive for eventual and germane PrE differentiation^7,9,25,62^. Interestingly, it has been reported ICM cell fate specification is maximally sensitive to exogenous FGF4 supplementation (promoting PrE differentiation) during early blastocyst maturation, with peak MEK/ERK inhibition sensitivity (preventing PrE differentiation) consistently observed at subsequent stages^9^. Although coincidental, the overlap in developmental timing of maximal FGF4 sensitivity and the requirement for p38-MAPK activity associated with PrE differentiation during early blastocyst maturation, may reflect a functional relationship. Indeed, we have previously reported that deficits in late blastocyst PrE cell numbers caused by sole inhibition of FGFR can be reversed by activating endogenous levels of p38-MAPK (via global over-expression of a constitutively active MKK6, one of two direct p38-MAPK activating kinases; an effect that can be blocked by simultaneous p38-MAPK chemical inhibition)^26^. Hence, it raises the possibility that FGF4 based regulation of PrE differentiation may have a component that is related to p38-MAPK activity (*i.e.* at those early blastocyst stages that precede the emergence of the mid-blastocyst salt & pepper pattern of specified EPI and PrE progenitors^10^). Theoretically this could include priming PrE progenitors for subsequent differentiation and/or directly facilitating necessary FGF-FGFR signalling (significantly we observed robustly reduced *Fgfr1* and *Fgfr2* mRNA expression, peaking at the E3.5 +7h time-point, after p38-MAPKi in our mRNA-Seq dataset; supplementary table S3a (+7h)) and would be consistent with global mobilisation of the translational apparatus (shown here to be impaired upon p38-MAPKi). However, a direct functional link between FGF4 signalling and p38-MAPK regulated translation remains to be demonstrated.

Notwithstanding these caveats, the central finding that p38-MAPK inhibition during early blastocyst maturation is associated with profound changes in both the (phospho)proteome and transcriptome of translation related genes (plus enhanced mRNA levels for genes associated with mitochondrial respiration – Figs. 2 & 3) and overall reduced levels of translation, transcription and rRNA processing (Fig. 5) is consistent with a role in creating metabolically permissive states compatible with PrE specification and differentiation. It is therefore significant that p38-MAPKi mediated PrE deficits can be partially restored by concomitant activation of mTOR (the well characterised metabolic regulator, the activity of which promotes active translation^47^ – Fig. 6f & j-q) and emphasises the importance of p38-MAPK mediated regulation of translation, and presumably the derived proteins, as a mechanism of facilitating PrE formation. The mTOR signalling pathway (integrated by two mTOR containing complexes, mTORC1 and mTORC2) is a global regulator of translation, and maintains cellular energy homeostasis in response to extra-cellular stimuli such as growth factors and nutrient availability^47^. As part of mTORC1, mTOR specifically controls transcription and translation of ribosomal proteins, synthesis and processing of rRNAs and ribosome biogenesis, assembly and availability, that ultimately dictates general translation of all mRNA transcripts (mTORC2 is associated with co-translational protein degradation and lipolysis)^47,63–66^. It was recently demonstrated that a prolonged state of *in vitro* developmental diapause (lasting 9-12 days) can be chemically induced in early stage (E3.5) mouse blastocysts upon dual inhibition of the mTORC1 and mTORC2, mimicking naturally occurring *in vivo* diapause^41^. The developmental diapause was associated with arrested cell division and reduced levels of global translation, and induced epigenetic chromatin remodelling causing reduced nascent transcription^41,42^; all phenotypes also observed after p38-MAPKi reported here (Fig. 5b-d, f & g). However, despite such similarities and the partial reversal of the PrE differentiation phenotype caused by simultaneous activation of mTOR (expressed as the emergent population of NANOG, GATA6 & GATA4 co-expressing ICM cells – Fig. 6p, q), it is clear from our results that p38-MAPKi is not a mere phenocopy of mTOR inhibition induced diapause. Rather, that p38-MAPKi is characterised by impaired PrE differentiation in the context of on-going development, that nevertheless has all the hallmarks of impaired translation. It is also notable that the restoration of GATA4 positive PrE cells in blastocysts co-treated with p38-MAPK inhibitor and mTOR activator was frequently accompanied by co-expression of both NANOG and GATA4 (Fig. 6f & q). This suggests such ICM cells represent a pseudo-state, not normally existing during unperturbed blastocyst maturation, whereby PrE differentiation has been uncoupled from the need to downregulate NANOG levels (as would be normal for PrE progenitors to exit their uncommitted state and specify PrE differentiation under ordinary developmental conditions). Interestingly, irrespective of co-treatment with mTOR inhibitor the number of EPI progenitors (defined by sole expression of NANOG) in p38-MAPKi blastocysts remained statistically unchanged (Fig. 6m). Not only does this again indicate EPI specification is not affected by p38-MAPKi, it also illustrates the specificity of the uncovered translational defects towards specification and differentiation of PrE progenitors. We also noted the partial PrE differentiation ‘rescue’ effects of mTOR activation on p38-MAPKi blastocyst were not associated with enhanced blastocyst cavity expansion (supplementary Fig. S3), suggesting these and the originally observed p38-MAPKi phenotypes are, at least partly, PrE progenitor cell autonomous. Furthermore, the partial nature of the observed rescue of PrE specification in blastocysts cultured under combined p38-MAPKi plus mTOR activated conditions, strongly suggest p38-MAPK mediated regulation will be multi-factorial. Notably, there is solid evidence from other systems of p38-MAPK mediated translational regulation acting via direct phosphorylation of MAPK-interacting kinases (MNKs); leading to phosphorylation of eIF4E and initiation of 5’-7-methylguanosine cap-dependent translation^46^. However, despite reports of reduced eIF4E phosphorylation in mouse genetic knockouts of *Mnk1* and/or *Mnk2,* no reductions in protein synthesis have been reported and full development of viable animals is sustained^67^. It will be of considerable future interest to precisely determine how p38-MAPK co-ordinately regulates early blastocyst translation in the context of PrE specification and differentiation.

The observed upregulation of ribosomal protein transcripts (peaking at the E3.5 +7h time-point, yet returning to control levels after 10 hours of p38-MAPKi – Fig. 3), that are also downregulated at the protein level (Fig. 2c & Fig. 3e), suggests their over-expression and/or accumulation is due to the overall impaired levels of translation we confirmed (Fig. 5b-d). In this context, it is important that we also noted a significant accumulation of unprocessed rRNA transcripts associated with p38-MAPKi (Fig. 5e), that could result (via a mechanism of diminished ribosome biogenesis) in the impaired translation levels observed (Fig. 2c & Fig. 5b-d). As referenced above, we were drawn to a study of HTATSF1, a factor known to regulate transcriptional and post-transcriptional processing of rRNA and mRNAs and thus, modulation of protein synthesis levels during the transition between pre- and post-implantation embryonic naïve and primed pluripotency, respectively^40^. HATASF1 mediated repression of active translation during this period was shown to be associated with increased intron retention in ribosomal protein encoding mRNAs, leading to their degradation by the nonsense mediated decay pathway^40^. Although we also observed reduced translation after p38-MAPKi (Fig. 2c & Fig. 5b-d) we did not find any evidence for similarly enhanced intron retention in our mRNA-Seq dataset (supplementary table S10). However, we did identify MYBPP1A in our phosphoproteome screen for candidate p38-MAPK effectors (Fig. 2e – the clonal downregulation of which is associated with reduced contribution to the late blastocyst stage PrE, Fig. 4), that was also confirmed as an interacting partner of HTATSF1^40^. Insights from other cell culture based studies implicate MYBBP1A in transcriptional repression of rRNA gene expression and as a facilitator of pre-rRNA transcript processing, with depletion of MYBBP1A resulting in the accumulation of unprocessed rRNA precursors^33^; a phenotype observed in p38-MAPKi blastocysts (Fig. 5e). Thus, we suggest it is possible MYBBP1A (under the direct/indirect regulation of p38-MAPK) may act as a checkpoint agent enabling switching from a homeostatic balance in cellular protein synthesis and the coordination of a functionally important translational response to externally provided differentiation cues (via regulation of mature rRNAs levels available for functional ribosome biogenesis). This hypothesis is strengthened by reports, in other models, describing MYBBP1A specific functions related to rRNA expression and processing during cellular stress response^35,36^. Additionally, MYBBP1A is also known to suppress mitochondrial respiration, via an interaction with the transcriptional coactivator PPARGC1A. Interestingly, our data have identified mitochondrial respiration related transcripts as the second most significant ontological class (after those related to translation) of p38-MAPKi mediated differentially expressed genes (Fig. 3d – representing upregulated transcript levels). In our phosphoproteomic screen, we detected multiple p38-MAPKi sensitive phosphorylation sites within the MYBBP1A protein (Fig. 2e & supplementary table S2d). Whereas phosphorylation at serine 1280 (S1280) decreased, we also detected increased phosphorylation at S1303 and S1323 among other sites. The only experimentally verified MYBBP1A phosphorylation site identified, to our knowledge, is centred on S1303 and is reported to be a substrate of the mitotic Aurora B kinase^68^. Our results indicate S1280 represents a good candidate to be a direct substrate of p38-MAPK, but this remains to be experimentally validated. Interestingly, we also identified p38-MAPKi mediated differential phosphorylation of PDCD11 (representing both enhanced and depleted phosphorylation at two independent sites – supplementary table S2d). Similar to MYBBP1A, PDCD11 has also been implicated in rRNA processing^69^, further suggesting p38-MAPK regulation of germane rRNA processing is an important factor during mouse blastocyst maturation. However, the mechanistic details of how such phosphorylation events functionally impact MYBBP1A (and/or PDCD11) and what extent they potentially play in the described early blastocyst p38-MAPKi induced PrE phenotype remain to be examined.

## Conclusion

In conclusion, we report a previously undefined and multi-layered role for p38-MAPK mediated translational regulation within a limited developmental window of early blastocyst maturation. p38-MAPK activity during this window ensures germane ICM cell fate specification, with specific reference to PrE derivation, by the late blastocyst (E4.5) stage. We propose such positive translational regulation is necessary to prime, or at least facilitate, subsequent differentiation of PrE progenitors (see model – Fig. 7).

**Figure 7:**
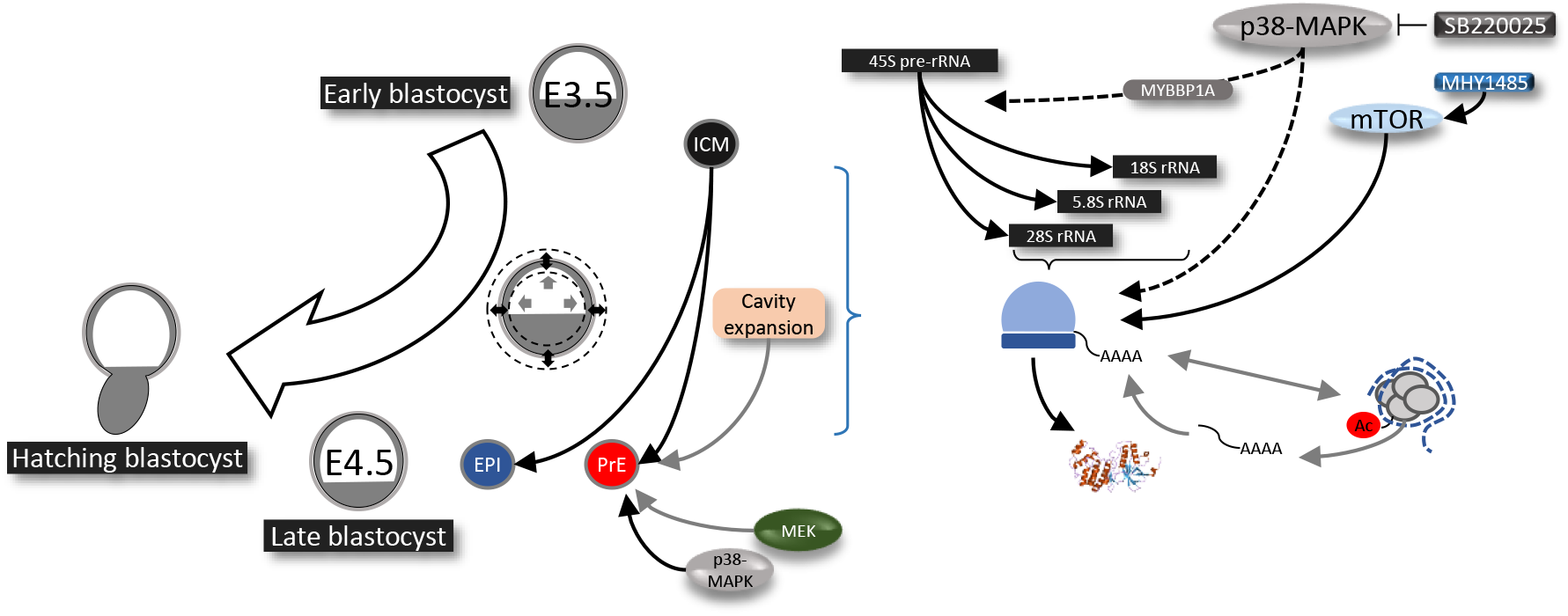
Working model of the role of p38-MAPK regulating protein translation in order to prime PrE differentiation during preimplantation stage blastocyst maturation.

## Methods

### Mouse lines and embryo culture

All mouse related experimental procedures (*i.e.* collecting preimplantation stage embryos for further study) complied with ‘ARRIVE’ guidelines and were carried out in accordance with EU directive 2010/63/EU (for animal experiments). Superovulation and strain mating regimen to produce experimental embryos were as follows; F1 hybrid (C57Bl6 x CBA/W) females were subject to peritoneal injection with 7.5 IU PMSG (Folligon^®^ MSD Animal Health) and 48 hours later with 7.5 IU hCG (Sigma-Aldrich^®^ cat. # CG10), followed by overnight F1 male mating (successful mating confirmed next morning by presence of vaginal plugs). E1.5 (*i.e.* 2-cell) stage embryos were isolated from the female oviducts in M2 media (prewarmed at 37°C for at least 2-3 hours) and thereafter cultured in KSOM (EmbryoMax^®^ KSOM Mouse Embryo Media; cat. # MR-020P-5F – pre-warmed and equilibrated in 5% CO2 and 37°C), either with or without amino acid (AA) supplementation. For KSOM+AA condition, Gibco™ MEM Non-Essential Amino Acids Solution (100X) (cat. # 11140035) and Gibco™ MEM Amino Acids Solution (50X) (cat. # 11130036) were used to a working concentration of 0.5X. Embryos were cultured in micro-drops prepared in 35mm tissue culture dishes covered with light mineral oil (Irvine Scientific. cat. # 9305), in 5% CO2 incubators maintained at 37°C until the appropriate stage. Pharmacological manipulations were carried out by addition of chemical agents dissolved in dimethyl sulfoxide (DMSO Sigma-Aldrich^®^ cat. # D4540) to KSOM±AA (concentration details *etc.,* in supplementary table S11). Equivalent volumes of solvent control were used to a final working concentration of 0.2-0.5% by volume. All KSOM based culture media, with or without additional chemicals (AAs, pharmacological agents), were pre-warmed and equilibrated in 5% CO2 and 37°C for at least 3-4 hours prior to embryo transfer.

### Sample collection for mass spectrometric analysing of differential (phospho)proteome

2-cell (E1.5) stage embryos were cultured in normal KSOM+AA conditions until E3.5 +2h and thereafter 300 embryos each were moved to control or p38-MAPKi conditions and cultured for another 7 hours (E3.5 +9h). The embryos were then washed through pre-warmed Hank’s balanced salt solution (HBSS; Sigma-Aldrich^®^ H9269) and lysed by moving to a 1.5ml centrifuge tube containing about 15μl of SDT-lysis buffer (4% (w/v) SDS, 100 mM Tris-HCl pH 7.6, 0.1 M DTT). Cell lysis was performed by incubating the tubes in a 95°C thermoblock for 12 minutes, brief centrifugation at 750 rpm, cooling to room temperature and storage at −80°C.

### Samples preparation for Liquid Chromatography-Mass Spectrometry (LC-MS) analyses

Individual protein solutions were processed by filter–aided sample preparation (FASP) method^70^ with some modifications. The samples were mixed with 8M UA buffer (8M urea in 100 mM Tris-HCl, pH 8.5), loaded onto the Microcon device with MWCO 30 kDa (Merck Millipore) and centrifuged at 7,000× g for 30 minutes at 20°C. The retained proteins were washed (all centrifugation steps after sample loading done at 14,000× g) with 200μl UA buffer. The final protein concentrates kept in the Microcon device were mixed with 100μl of UA buffer containing 50 mM iodoacetamide and incubated in the dark for 20 minutes. After the next centrifugation step, the samples were washed three times with 100μl UA buffer and three times with 100μl of 50 mM NaHCO3. Trypsin (1μg, sequencing grade, Promega) was added onto the filter and the mixture was incubated for 18 h at 37°C. The tryptic peptides were finally eluted by centrifugation followed by two additional elutions with 50μL of 50mM NaHCO3. Peptides were then cleaned by liquidliquid extraction (3 iterations) using water saturated ethyl acetate^71^.

1/10^th^ of the cleaned FASP eluate was taken out for direct LC-MS measurements, evaporated completely in SpeedVac concentrator (Thermo Fisher Scientific). Peptides were further transferred into LC-MS vials using 50μL of 2.5% formic acid (FA) in 50% acetonitrile (ACN) and 100μL of pure ACN and with addition of polyethylene glycol (final concentration 0.001%) and concentrated in a SpeedVac concentrator.

Phosphopeptides were enriched from the remaining 9/10^th^ of the cleaned FASP eluate after complete solvent evaporation (SpeedVac concentrator) using High-Select™ TiO2 Phosphopeptide Enrichment Kit (Thermo Fisher Scientific, Waltham, Massachusetts, USA) according to manufacturer protocol. Phosphopeptide standards (0.1 pmol of MS PhosphoMix 1, 2, 3 Light; Sigma-Aldrich, St. Louis, Missouri, USA) was added to suspended sample in binding/equilibration buffer. Flow-through fraction was dried and used for second enrichment step using High-Select™ Fe-NTA Phosphopeptide Enrichment Kit (Thermo Fisher Scientific, Waltham, Massachusetts, USA) according to manufacturer protocol. Resulting phosphopeptides were extracted into LC-MS vials by 2.5% FA in 50% ACN and 100% ACN with addition of polyethylene glycol (final concentration 0.001%) and concentrated in a SpeedVac concentrator).

### LC-MS analysis of peptides

LC-MS/MS analyses of all peptide mixtures (3 peptide solutions for each sample – 1) not enriched; 2) enriched on phosphopeptides using TiO2 enrichment kit and; 3) enriched on phosphopeptides using Fe-NTA enrichment kit) were done using RSLCnano system connected to Orbitrap Fusion Lumos mass spectrometer (Thermo Fisher Scientific). Prior to LC separation, tryptic digests were online concentrated and desalted using trapping column (100 μm × 30 mm, column compartment temperature of 40°C) filled with 3.5-μm X-Bridge BEH 130 C18 sorbent (Waters). After washing of trapping column with 0.1% FA, the peptides were eluted (flow rate – 300nl/min) from the trapping column onto an analytical column (Acclaim Pepmap100 C18, 3 μm particles, 75 μm × 500 mm; column compartment temperature of 40°C, Thermo Fisher Scientific) by using 50 or 100 minutes long nonlinear gradient program (1-56% of mobile phase B; mobile phase A: 0.1% FA in water; mobile phase B: 0.1% FA in 80% ACN) for analysis of phosphopeptide enriched fractions or not enriched peptide mixtures, respectively. Equilibration of the trapping column and the analytical column was done prior to sample injection to sample loop. The analytical column outlet was directly connected to the Digital PicoView 550 (New Objective) ion source with sheath gas option and SilicaTip emitter (New Objective; FS360-20-15-N-20-C12) utilization. ABIRD (Active Background Ion Reduction Device, ESI Source Solutions) was installed.

MS data were acquired in a data-dependent strategy with cycle time for 3 seconds and with survey scan (350-2000 m/z). The resolution of the survey scan was 60000 (200 m/z) with a target value of 4×10^5^ ions and maximum injection time of 50ms. HCD MS/MS (30% relative fragmentation energy, normal mass range) spectra were acquired with a target value of 5.0×10^4^. The MS/MS spectra were recorded in Orbitrap at resolving power of 30,000 or 15,000 (200 m/z) and the maximum injection time for MS/MS was 500 or 22 ms for analysis of phosphopeptides enriched fraction or non-enriched peptide mixture, respectively. Dynamic exclusion was enabled for 60 seconds after one MS/MS spectra acquisition. The isolation window for MS/MS fragmentation was set to 1.6 m/z.

The analysis of the mass spectrometric RAW data files was carried out using the MaxQuant software (version 1.6.6.0) using default settings unless otherwise noted. MS/MS ion searches were done against modified cRAP database (based on http://www.thegpm.org/crap, 112 protein sequences) containing protein contaminants like keratin, trypsin etc., and UniProtKB protein database for *Mus musculus* (ftp://ftp.uniprot.org/pub/databases/uniprot/current_release/knowledgebase/reference_proteomes/Eukaryota/UP000000589_10090.fasta.gz; downloaded 8.5.2019, version 2019/05, number of protein sequences: 22,287). Oxidation of methionine and proline, deamidation (N, Q) and acetylation (protein N-terminus) as optional modification, and trypsin/P enzyme with 2 allowed miss cleavages and minimal peptide length 6 amino acids were set. Peptides and proteins with FDR threshold <0.01 and proteins having at least one unique or razor peptide were considered only. Match between runs was set separately for all enriched and not enriched peptide solution analyses. Experiment name was set the same for differently enriched peptide solution analyses coming from the same sample.

Protein intensities reported in proteinGroups.txt file and evidence intensities reported in evidence.txt file (output of MaxQuant program) were further processed using the software container environment (https://github.com/OmicsWorkflows), version 3.7.1a. Processing workflow is available upon request. Briefly, it covered: 1) protein level data processing: a) removal of decoy hits and contaminant protein groups, b) protein group intensities log2 transformation, c) LoessF normalization and d) differential expression using LIMMA statistical test (qualitative changes were considered separately without statistical evaluation); 2) phosphopeptide level: a) removal of not phosphorylated and contaminant proteins associated evidences, b) grouping of the evidences for the identical combination of peptide sequence and set of modifications, c) LoessF normalization and d) differential expression using LIMMA statistical test (qualitative changes were considered separately without statistical evaluation).

Proteins and phosphopeptide candidates were selected based on the criteria as mentioned within the results.

### mRNA-Seq library preparation and sequencing

2-cell (E1.5) stage embryos were cultured in normal KSOM+AA conditions until E3.5 and thereafter 30 embryos each were moved to control or p38-MAPKi conditions and cultured for another 4, 7 or 10 hours. The embryos were then processed for total RNA isolation using the ARCTURUS^®^ PicoPure^®^ RNA Isolation Kit (Applied Biosystems™; KIT0204), following the manufacturer’s protocol. Oligo dT-based mRNA isolation was carried out using the NEBNext^®^ Poly(A) mRNA Magnetic Isolation Module (NEB; E7490) and libraries were prepared using the NEBNext^®^ Ultra™ II Directional RNA Library Prep Kit for Illumina^®^ (NEB; E7760), as directed by the manufacturer. Libraries were sequenced as 50 bp single-end reads on the Illumina^®^ HiSeq4000 platform. RNA-seq datasets are available in GEO database with accession number GSE162233.

### Data processing and bioinformatics analysis

Sequenced reads were adapter- and quality-trimmed using Trim Galore! v0.4.1 and mapped to the mouse GRCm38 genome with Hisat2 v2.0.5. Differentially expressed genes between control and p38-MAPKi embryos at individual time-points were identified by DESeq2 within Seqmonk v1.45.4 using GRCm38_v90 transcriptome annotation. Hierarchical clustering was performed in Seqmonk v1.45.4 with mean-centred expression values. Gene ontology was performed using GOrilla^72,73^.

### Embryo manipulation by microinjections

Single (for immuno-fluorescence confocal microscopy) or double (for qRT-PCR) blastomere microinjections of 2-cell (E1.5) stage embryos were performed using FemtoJet or FemtoJet 4i (Eppendorf; 5252000013) micro-injectors, mechanical micro-manipulators (Leica; ST0036714) and CellTram Vario (Eppendorf; 5176000033) pneumatic handler, under a negative capacitance enabled current controlled by an Electro 705 Electrometer (WPI; SYS-705) and on the stage of an Olympus IX71 inverted fluorescence microscope. Embryos were pneumatically handled and immobilised for microinjection using a borosilicate glass capillary holder (without filament – Harvard Apparatus; 30-0017). Micro-injectors were connected to needles prepared from filamented borosilicate glass capillaries (Harvard Apparatus; 30-0038) using a Narishige PC-10 capillary glass needle puller. All siRNAs (supplementary table S12) were co-microinjected at 10μM concentrations, with 50ng/μl *H2b-RFP* mRNA, in pre-warmed drops of M2 media overlaid with mineral oil, on the surface of concaved microscope slides.

### Bright-field microscopy, immunofluorescence staining, confocal microscopy and image acquisition

Bright-field images were captured using an Olympus IX71 inverted fluorescence microscope and Optika TCB3.0 imaging unit, plus associated Optika Vision Lite 2.1 software. To remove the *zona pellucida*, blastocysts were quickly washed and pipetted in pre-warmed drops of acidic Tyrode’s Solution (Sigma-Aldrich^®^ cat. # T1788) until visually undetectable, immediately followed by washes through pre-warmed drops of M2 media. Thereafter embryos were fixed, in the dark, with 4% paraformaldehyde (Santa Cruz Biotechnology, Inc. cat. # sc-281692) for 20 minutes at room temperature. Permeabilisation was performed by transferring embryos to a 0.5% solution of Triton X-100 (Sigma-Aldrich^®^ cat. # T8787), in phosphate buffered saline (PBS), for 20 minutes at room temperature. All washes post-fixation, permeabilisation and antibody staining were performed in PBS with 0.05% TWEEN^®^ 20 (Sigma-Aldrich^®^ cat. # P9416) (PBST) by transferring embryos between two drops or wells (of 96-well micro-titre plates) of PBST, for 20 minutes at room temperature. Blocking and antibody staining was performed in 3% bovine serum albumin (BSA; Sigma-Aldrich^®^ cat. # A7906) in PBST. Blocking incubations of 30 minutes at 4°C were performed before both primary and secondary antibody staining; primary antibody staining (in blocking buffer) was incubated overnight (~16 hours) at 4°C and secondary antibody staining carried out in the dark at room temperature for 70 minutes. Stained embryos were mounted in DAPI containing mounting medium VECTASHIELD^®^ (Vector Laboratories, Inc. cat. # H-1200), placed under cover slips on glass-bottomed 35mm culture plates and incubated at 4°C for 30 minutes in the dark, prior to confocal imaging. Details of the primary and secondary antibody combinations used can be found in supplementary table S13. Confocal images were acquired using a FV10i Confocal Laser Scanning Microscope and FV10i-SW image acquisition software (Olympus^®^). Images were analysed using FV10-ASW 4.2 Viewer (Olympus^®^) and Imaris X64 Microscopy Image Analysis Software (version 6.2.1; Bitplane AG – Oxford Instruments plc). Cells were counted both manually and semi-automatically using Imaris X64.

### Time-lapse imaging

Embryos at E3.5 were placed in 20μl droplets of KSOM+AA (+DMSO or +SB220025) on 16-well dishes (PrimoVision, Vitrolife), one embryo per well, within a PrimoVision imaging system and cultured for 24 hours (until E4.5) under standard culture conditions. The embryos were imaged every 10 minutes and the equatorial area was recorded and data analysed (Fig. 1c and as described previously^74^).

### Scarce sample polysome (SSP) profiling

2-cell (E1.5) stage embryos were cultured in KSOM+AA and transferred to control or p38-MAPKi conditions for 10 hours starting from E3.5 (E3.5 +10h; as depicted in Fig. 5a). Prior to harvesting, the embryos were transferred to cycloheximide (0.1mg/ml) containing media for 10 minutes. This was followed by 3x washes in PBS (supplemented with 0.1mg/ml cycloheximide) and 1x wash with transfer medium (0.1% PVA in PBS + 0.1mg/ml cyclohexamide). The embryos were then transferred to 0.5ml DNA low-bind tubes in a minimal volume of transfer medium, flash frozen in liquid nitrogen and stored at −80°C. Cellular lysis and sucrose gradient centrifugation to procure ribosomal fractions, followed by isolation of corresponding ribosomal RNA was performed as described previously^39^. The resultant fractions representing rRNA corresponding to non-polysomes (fractions 1 to 5; from the upper *i.e.* lighter fractions) and polysomes (fractions 6 to 10; from the lower *i.e.* heavier fractions) were resuspended in 20μl of nuclease-free water. cDNA was then prepared as follows: 4.0μl of total RNA was mixed with 1.6μl of 0.183μg/μl random hexamers and 3.0μl of nuclease-free water. The samples were incubated on a thermoblock at 70°C for 5 minutes followed by ice for 5 minutes and were then briefly microfuged. 3.9μl of nuclease-free water, 4.0μl of 5x reaction buffer, 1.0μl of RevertAid H Minus reverse transcriptase (Thermo Scientific™; EP0451), 0.5μl of RiboLock RNase inhibitor (Thermo Scientific™; EO0381) and 2.0μl of dNTP mix (Thermo Scientific™; R0192) were then added. The preparations were processed in a thermocycler with the following program: 25°C for 10 minutes, 37°C for 5 minutes, 42°C for 75 minutes and 70°C for 10 minutes. Synthesised cDNA was stored at −20°C and 2.0μl aliquots used as template for each qRT-PCR reaction (as described previously^39^).

### O-propargyl-puromycin (OPP) quantification of *de novo* protein synthesis

Mouse embryos, *in vitro* cultured from E1.5 to E3.5 in KSOM+AA, and then further cultured under control or p38-MAPKi conditions for 10 hours (E3.5 +10h). OPP staining was then performed using the Click-iT^®^ Plus OPP Alexa Fluor^®^ 488 Protein Synthesis Assay Kit (Invitrogen™; C10456), following the prescribed protocol modified for *in vitro* preimplantation embryo culture. In brief, equilibrated culture dishes (described above) containing drops of KSOM (without amino acid supplementation) and comprising 40μM solutions of OPP and supplemented with either control DMSO or 20μM SB220025 were prepared.

Following a 60 minute pre-incubation period, control and p38-MAPKi blastocysts were transferred to their respective OPP containing dishes and incubated for a further 60 minutes. Thereafter embryos were fixed, permeabilised and washed as described in the above section relating to immunofluorescence confocal microscopy. Click-iT^®^ reactions, to enable fluorescent visualisation of OPP into nascent polypeptides, were set up as described in the kit (comprising Click-iT^®^ reaction buffer, copper protectant, Alexa Fluor^®^ 488 picolyl azide and reaction buffer additive) and the fixed embryos were incubated in reaction mixture for 25 minutes at room temperature, in the dark. Thereafter, embryos were washed in a 1:1 mixture of kit provided wash buffer and PBST, mounted in VECTASHIELD^®^ and imaged using standard confocal microscope settings (as described above) to detect the Alexa Fluor^®^ 488 fluorophore and quantified as described.

### Cell number quantification, statistics and graphical representation

Total embryo cell number counts (plus outer and inner cell populations) from confocal acquired micrographs (based on DAPI nuclei staining) were further subcategorised as EPI or PrE cells based on detectable and exclusive expression NANOG and GATA4 protein expression (confocal images in Fig. 1a, 4 & 6 and graphs in Fig. 1a, 4 & 6). Cells not located within blastocyst ICMs that also did not stain for either GATA4 and/or NANOG, were designated as outer/TE cells. In relation to Fig. 6 & 6p specifically, ICM cells that were positively stained for both GATA6 and NANOG at E4.5 were designated as uncommitted/unspecified in terms of cell fate and the population of similarly GATA4 and NANOG positive ICM cells (primarily resulting from co-inhibition/co-activation of p38-MAPK and mTOR, respectively – Fig. 6f) also recorded. In figures 5f and g, cells were designated as “outer cells” based on detectable CDX2 (TE marker^44^) expression. Relative levels of H4K16ac modified chromatin (Fig. 5f) were quantified to denote global chromatin status and quantified RNA polymerase-II S2p (Fig. 5g) levels to denote active transcriptional elongation, as described below (fluorescence intensity quantification and statistical analysis). Initial recording and data accumulation was carried out using Microsoft Excel and further statistical analysis and graphical representations were performed with GraphPad Prism 8. Based on the normality and lognormality comparisons, appropriate statistical tests were used for the compared datasets (summarised in supplementary table S14). Unless otherwise stated within individual graphs as a specific P value (if statistically insignificant), the stated significance intervals were depicted as follows: P value < 0.0001 (****), 0.0001 to 0.001 (***), 0.001 to 0.01 (**) and 0.01 to 0.05 (*). All graphs represent dot plots of the total sample size, with associated means and standard deviation bars provided.

### Fluorescence intensity quantification and statistical analysis

Fluorescence intensity pertaining to OPP incorporation (Fig. 5b) was quantified for the minimal complete confocal z-series of each embryo as a whole using Fiji (ImageJ)^75^. The confocal images from the channel pertaining to the 488nm wavelength were analysed as such: Image>Stacks>Z Project. The Projection type “Sum Slices” was used to obtain a Z-projected image for the entire embryo. Levels of H4K16ac (Fig. 5f) and RNA polymerase-II S2p (Fig. 5g) were quantified as fluorescence intensity per cell and differentiated into inner and outer cells based on absence or presence of CDX2 immuno-fluorescence, respectively. The measurements settings for all of the above were as follows: Analyze>Set Measurements; and the following options were chosen: Area, Mean grey value and Integrated density. Using the “Polygon selections” tool, an area encompassing the Z-projected embryo image (for OPP) or individual cell nuclei (for H4K16ac and Pol II S2p) were demarcated and the aforementioned measurements recorded. The selected area was then moved so as to encompass an area excluding that of the embryo or cell nucleus, respectively, and background measurements were recorded. This process was continued for all the embryos analysed, under both control and p38-MAPKi conditions and the results were transferred to a spreadsheet for further calculations. The Corrected Total Cell Fluorescence (CTCF), in arbitrary units, for each embryo was measured as such: CTCF = Integrated Density – (Area of selected cell X Mean fluorescence of background readings)^76,77^ – supplementary tables 5a-c. The calculated CTCF are plotted as scatters, with mean and standard deviations marked. The CTCF differences were statistically tested using Mann-Whitney test and the results, unless otherwise stated within individual graphs as a specific P value (if statistically insignificant), are stated as following significance intervals: P value < 0.0001 (****), 0.0001 to 0.001 (***), 0.001 to 0.01 (**) and 0.01 to 0.05 (*).

### Blastocyst size and volume calculations

Equatorial areas of blastocoel cavities (Fig. 1b) were calculated by measuring the inner circumference of the centrally located widest Z-stack using Fiji (ImageJ)^75^. In order to compare the volume of the p38-MAPK inhibited blastocoel cavities with that of the ones reported in Ryan et al.^12^, we deduced the volume from measurements of the inner circumference as well. Calculated area and corresponding volume measurements are tabulated in supplementary table S1c. Equatorial area and total volume measurements for the blastocyst as a whole was carried out by similarly measuring the outer circumference (as reported in supplementary Fig. 3). The measurements were set as follows: Analyze>Set Measurements; and the “Perimeter” option was selected; using the “Polygon selections” tool, the inner circumference was traced and measured. The radius of the measured circumference was deduced and that value was used to calculate an approximate equatorial section area (or volume) for all the embryos analysed under both control and p38-MAPKi conditions. The calculated area in μm^2^ (or volume in picoliters (pL)) is plotted as a scatter, with mean and standard deviations marked. The differences were statistically tested using Mann-Whitney test and the results, unless otherwise stated within individual graphs as a specific P value (if statistically insignificant), are stated at the following significance intervals: P value < 0.0001 (****), 0.0001 to 0.001 (***), 0.001 to 0.01 (**) and 0.01 to 0.05 (*).

### Quantitative real-time PCR (qRT-PCR)

Embryos from each stated experimental condition were collected and immediately processed for RNA extraction and isolation using the ARCTURUS^®^ PicoPure^®^ RNA Isolation Kit (Applied Biosystems™; KIT0204), following the manufacturer’s protocol. The entire eluted volume of total RNA was immediately DNase treated with TURBO™ DNase (Invitrogen™; AM2238) according to the manufacturer provided protocol. The whole sample was then subject to cDNA synthesis using SuperScript™ III Reverse Transcriptase (Invitrogen™; 18080044), as directed by the manufacturer and employing oligo d(T)16 (for Fig. 1d & 4b) (Invitrogen™; N8080128) or random hexamers (for Fig. 5c, d & e) (Thermo Scientific™; SO142), dNTP Mix (ThermoScientific™; R0192) and RNase Inhibitor (Applied Biosystems™; N8080119). The synthesized cDNA was diluted as required with nuclease free water and 1μl was used in 10μl individual final SYBR-green based qRT-PCR reaction volumes (qPCRBIO SyGreen Mix Lo-ROX; PB20.11). A Bio-Rad CFX96 Touch Real-Time PCR Detection System apparatus, employing standard settings, was employed for data accumulation and initial analysis was performed with the accompanying Bio-Rad CFX Manager™ software. Triplicate measurements per gene (the sequence of individual oligonucleotide primers, used at a final concentration of 300nM, are provided in supplementary table S15) were assayed from two biological replicates that were each technically replicated. The averaged transcript levels of analysed genes were derived after internal normalisation against *Tbp* or *H2afz* mRNA levels. As such, data was acquired and initially analysed with CFX Manager™, then processed in Microsoft Excel (biological and technical replicate averaging) and GraphPad Prism 8 (graphical output). Statistical tests used are detailed within supplementary table S14. Unless otherwise stated within individual graphs as a specific P value (if statistically insignificant), the stated significance intervals were depicted as: P value < 0.0001 (****), 0.0001 to 0.001 (***), 0.001 to 0.01 (**) and 0.01 to 0.05 (*); error bars denote calculated standard deviations.

## Supporting information

Single control embryo

Control embryos replicate 1

Control embryos replicate 2

Single p38-MAPK inhibited embryo

p38-MAPK inhibited embryos replicate 1

p38-MAPK inhibited embryos replicate 2

## Contributions

P.B. and A. W. B. conceived the project, designed experiments, analysed results and wrote the manuscript. P.B. conducted experiments. P.B. and L.G. prepared samples for mass spectrometry and RNA sequencing. L.G. analysed proteomic, phosphoproteomic and transcriptomic data and wrote associated portions within the manuscript. T.M. and M.P. designed and produced SSP profiling fractions. A.H. analysed images for blastocoel cavity volume. D.P. and Z.Z. performed phosphoproteomic mass spectrometry, did preliminary data analysis and wrote the associated methods. A.A. performed time-lapse imaging and analysis. D.J. and A.S. performed peripheral experiments.

## Availability of data and materials

RNA-seq datasets are available in GEO database with accession number GSE162233. Mass spectrometry datasets are available on request.

## Acknowledgments

We acknowledge the Institute of Parasitology (Biology Centre of the Czech Academy of Sciences, in České Budějovice) for housing mice, Marta Gajewska (Institute of Oncology, Warsaw, Poland) and Anna Piliszek (Institute of Genetics and Animal Breeding, Polish Academy of Sciences, Jastrzębiec, Poland) for founder CBA/W mice, Alena Krejčí (Faculty of Science, University of South Bohemia, Czech Republic) for pooling resources and other members of our laboratory for valuable inputs and discussions. We further acknowledge Markéta Hančová (IAPG, Liběchov) for her help in carrying out experiments. We also acknowledge the core facility Masaryk University, Brno, Czech Republic – Faculty of Informatics, supported by the MEYS CR (LM2018129 Czech-BioImaging) for assistance with image analysis. CIISB, Instruct-CZ Centre of Instruct-ERIC EU consortium, funded by MEYS CR infrastructure project LM2018127, is gratefully acknowledged for the financial support of the measurements at the CEITEC Proteomics Core Facility. Computational resources were supplied by the project “e-Infrastruktura CZ” (e-INFRA LM2018140) provided within the program Projects of Large Research, Development and Innovations Infrastructures.

