## Supplementary figures for "p38-MAPK mediated rRNA processing and translation regulation enables PrE differentiation during mouse blastocyst maturation"

Supplementary Fig. S1

a) Genes significantly differentially expressed in E3.5 +4 hours.

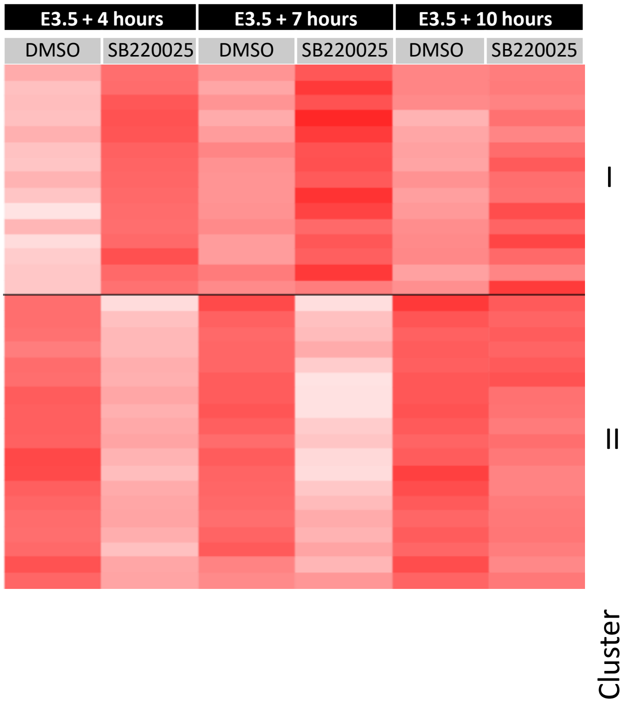


b) Genes significantly differentially expressed in E3.5 +10 hours.

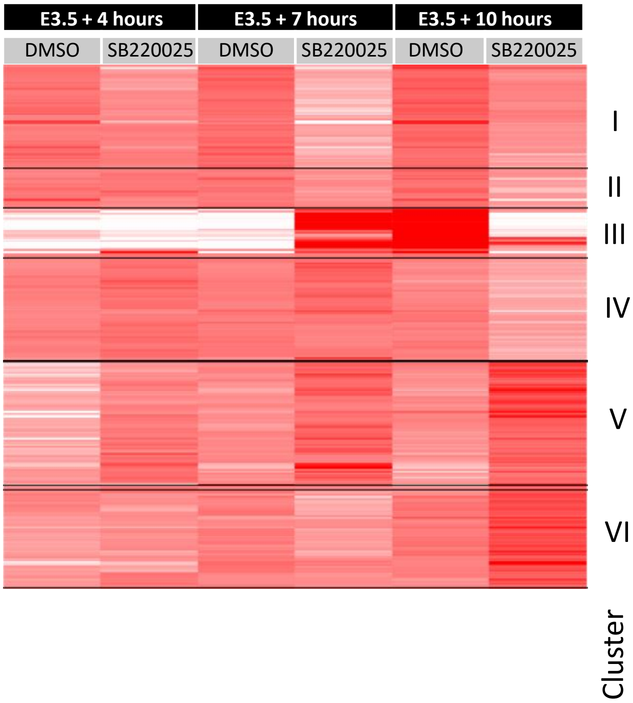


Hierarchical clustering heat-map depicting the expression of significantly changing gene mRNAs elicited by p38-MAPKi at the +4h (a) and +10h (b) time-points and the status of the transcript levels of those same genes at the two other time-points; forming two and six distinct expression clusters respectively (details in supplementary tables S7a and S7b respectively).

Supplementary Fig. S2

Embryo survival curve E3.5 onwards

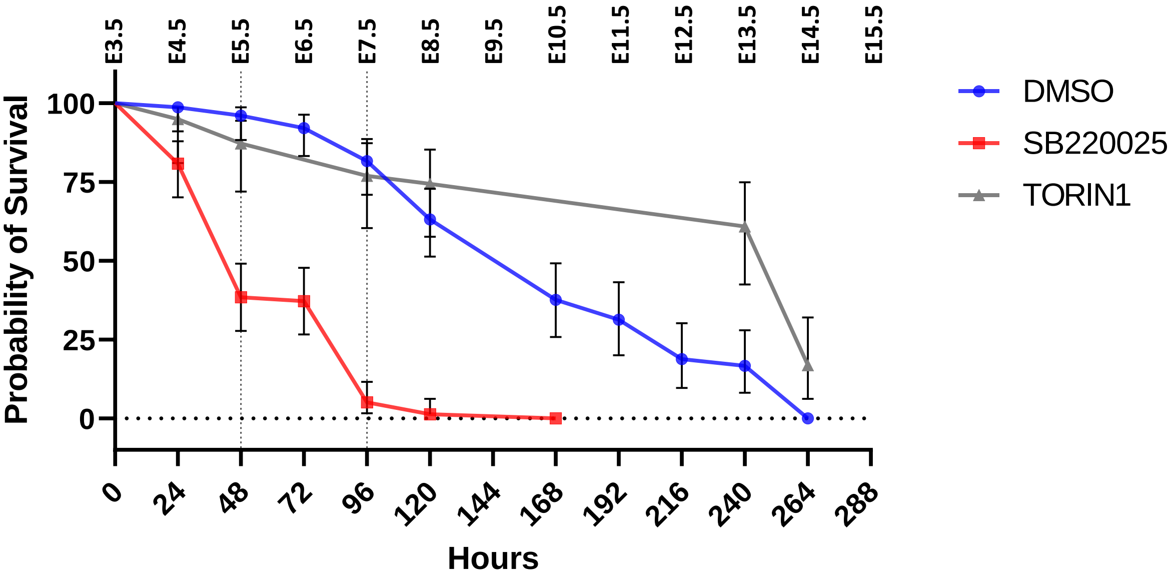


Brightfield images based analysis of survival of embryos cultured in control (DMSO; n=76), p38-MAPK inhibited (SB220025; n=78) and mTOR inhibited (TORIN1; n=39) conditions starting from E3.5.

Supplementary Fig. S3

Blastocyst size for embryos inhibited for 24 hours (E3.5 to E4.5)

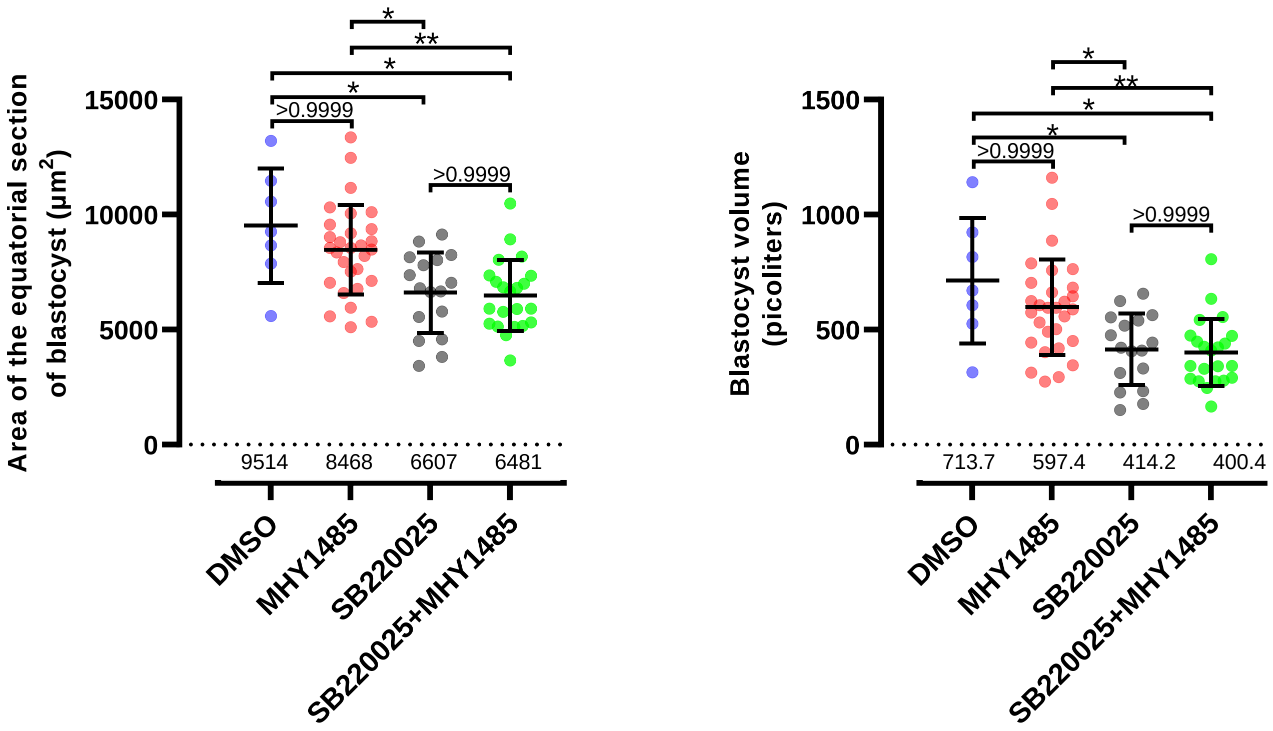


Quantification of blastocyst equatorial area (µm$^2$) and volume (picoliters, pL) in fixed blastocysts cultured (E3.5-E4.5) in control (DMSO; n=7), mTORa (MHY1485; n=29), p38-MAPKi (SB220025; n=17) and p38-MAPKi + mTORa (SB220025+MHY1485; n=22) conditions.
